# TEAD-targeting small molecules induce a cofactor switch to regulate the Hippo pathway

**DOI:** 10.1101/2024.11.15.623512

**Authors:** Alissa D. Guarnaccia, Thijs J. Hagenbeek, Wendy Lee, Noelyn Kljavin, Meena Choi, Gözde Ulas, Vasumathi Kameswaran, Daniel Le, Sayantanee Paul, Samir Vaidya, Jason R. Zbieg, James J. Crawford, Bence Daniel, Anwesha Dey, Jennie R. Lill

## Abstract

TEAD proteins are the main transcriptional effectors of the Hippo signaling pathway and a pharmacological target in oncology. Most TEAD-targeting small molecules act by disrupting interaction with the oncogenic transcriptional activators YAP and TAZ. Here, we describe an alternative mechanism for TEAD lipid pocket binding molecules. We report that select sulfonamide-containing compounds promote TEAD interaction with the transcriptional repressor VGLL4 to induce a small molecule-mediated cofactor switch from YAP to VGLL4. Chemically induced VGLL4–TEAD complexes counteract YAP activity at chromatin to repress pro-growth gene networks, including genes involved in cellular proliferation and mechanosignaling. VGLL4 is required for an anti-proliferative response to these select compounds, and genetic deletion of VGLL4 causes resistance to these molecules *in vitro* and *in vivo*. Our data reveal a category of molecules that facilitate the repressive VGLL4–TEAD interaction and open up new understandings for curbing the oncogenic activity of Hippo pathway deregulation.

## INTRODUCTION

Initially discovered in *Drosophila*, the Hippo pathway is now realized as a key regulatory mechanism in human development and disease, including cancer.^1^ The Hippo signaling pathway is a crucial cellular regulator, exerting tumor suppressive activity to tightly control cellular growth, proliferation, and differentiation.^2^ The core activity of the Hippo pathway involves upstream factors, including MST1/2, NF2, and LATS1/2, which promote phosphorylation of YAP and TAZ to prevent their nuclear translocation and transcriptional activity with the TEAD family of transcription factors (TEAD1-4). In cancer, deregulation of the Hippo pathway is prominent both as a driver of tumor development (for example NF2 mutations ^3–5^ and inactivation of LATS kinases ^6,7^) and as a mechanism of acquired resistance to targeted therapies.^8–13^ Therefore, modalities that can effectively reactivate the tumor-suppressive function of the Hippo pathway hold significant therapeutic potential.

Understanding how the Hippo pathway functions is essential to targeting it effectively. The key output of the Hippo pathway is transcriptional regulation mediated by TEAD transcription factors. Because TEAD proteins have minimal inherent transcriptional activity, TEAD relies on interactions with other proteins in order to influence chromatin. Interaction partners include other transcription factors,^14–16^ chromatin remodelers,^17^ and DNA damage repair proteins,^18,19^ as well as coactivators and corepressors. In particular, TEAD coactivators and corepressors do not have DNA binding domains and thus depend on direct interaction with TEAD in order to engage with chromatin. The three TEAD cofactors YAP, TAZ, and VGLL4 are notable because they have mutually exclusive, overlapping binding sites on TEAD ^20^ and engage TEAD to influence a transcriptional balancing act of activation and repression.^21–23^ YAP and TAZ are TEAD coactivators ^1,2,24^ and can be oncogenic, as they are frequently observed to have high nuclear levels and activity in cancers.^8,24,25^ In contrast, VGLL4 is a TEAD corepressor and is tumor suppressive,^23,26–28^ and low expression of *VGLL4* is associated with poor prognosis in a variety of malignancies.^21,22,29–32^ Proof-of-concept experiments using VGLL4-mimicking peptides have demonstrated that leveraging the VGLL4–TEAD interaction can counteract YAP to suppress tumor growth.^22,33^ However, the extent to which VGLL4–TEAD can be modulated by drug-like small molecules remains largely unknown.

TEAD small molecule inhibitors represent the most prominent strategy for targeting deregulated Hippo signaling in cancer.^20,34^ Unlike YAP and TAZ which are intrinsically disordered proteins, TEAD proteins contain a druggable hydrophobic pocket.^35,36^ This site is termed the lipid pocket because it is post-translationally occupied by a lipid modification, *S*-palmitoylation of a conserved cysteine.^37,38^ Small molecules are designed to specifically engage this pocket with the objective of allosterically disrupting the YAP–TEAD interaction to repress oncogenic transcriptional programs.^20,34,35,39^ Development of TEAD lipid pocket-targeting therapeutic candidates is advancing rapidly, with only six years between the discovery of this druggable pocket ^35,36,40^ and the entrance of such molecules in the clinic (NCT05228015, NCT04665206). With this rapid advancement, deeper biological insight is needed to understand precisely how chemical engagement at the lipid pocket modulates TEAD, particularly regarding the impact of inhibitory molecules on TEAD protein interactions. Although many TEAD-targeting molecules potently inhibit the YAP–TEAD interaction,^8,34,41,42^ paradoxically, some molecules inhibit TEAD palmitoylation and repress Hippo target gene expression without disrupting the YAP–TEAD interaction.^34,43–45^ If disruption of YAP is not necessary for efficacy of TEAD inhibitors, other mechanistic forces are likely at play.

Here, we challenge the idea that TEAD lipid pocket binding (LPB) inhibitors act exclusively by directly disrupting the YAP/TAZ–TEAD interaction. We take a quantitative, global proteomic approach to interrogate how TEAD interactions change upon LPB inhibitor treatment. Our findings delineate a class of TEAD-targeting molecules that boosts the VGLL4–TEAD interaction and provide insight into the repressive transcriptional effects of VGLL4. We show that, distinct from direct YAP/TAZ displacement, an enhanced VGLL4–TEAD axis can counteract YAP oncogenic activity and is an alternative mechanism of anti-proliferative action for TEAD LPB inhibitors. Altogether, we report on key details of the balance of protein interactions that govern TEAD transcriptional activity, and we demonstrate how small molecules can induce a cofactor switch to tune this balance.

## RESULTS

### Lipid pocket binding compound, Compound 2, increases the interaction of TEAD and VGLL4

To better understand the cellular effects of TEAD LPB inhibitors, we used the Hippo-driven mesothelioma cell line NCI-H226, a workhorse model for TEAD studies.^8,41,42,46–49^ We focused on two characterized pan-TEAD LPB small molecules: GNE-7883 (lipid pocket affinity of ∼330 nM)^8^ and Compound 2 (lipid pocket affinity of ∼20 nM)^43^ (Figure 1A). We chose these molecules because they differ in their impact on the YAP/TAZ–TEAD interaction: GNE-7883 allosterically blocks the interaction between TEAD and YAP/TAZ,^8^ while Compound 2 does not.^43^ Despite the difference, both molecules decrease the cellular viability of NCI-H226 cancer cells with nanomolar EC_50_ values (Figure 1B), similar to what has previously been reported.^8,43^

**Figure 1:**
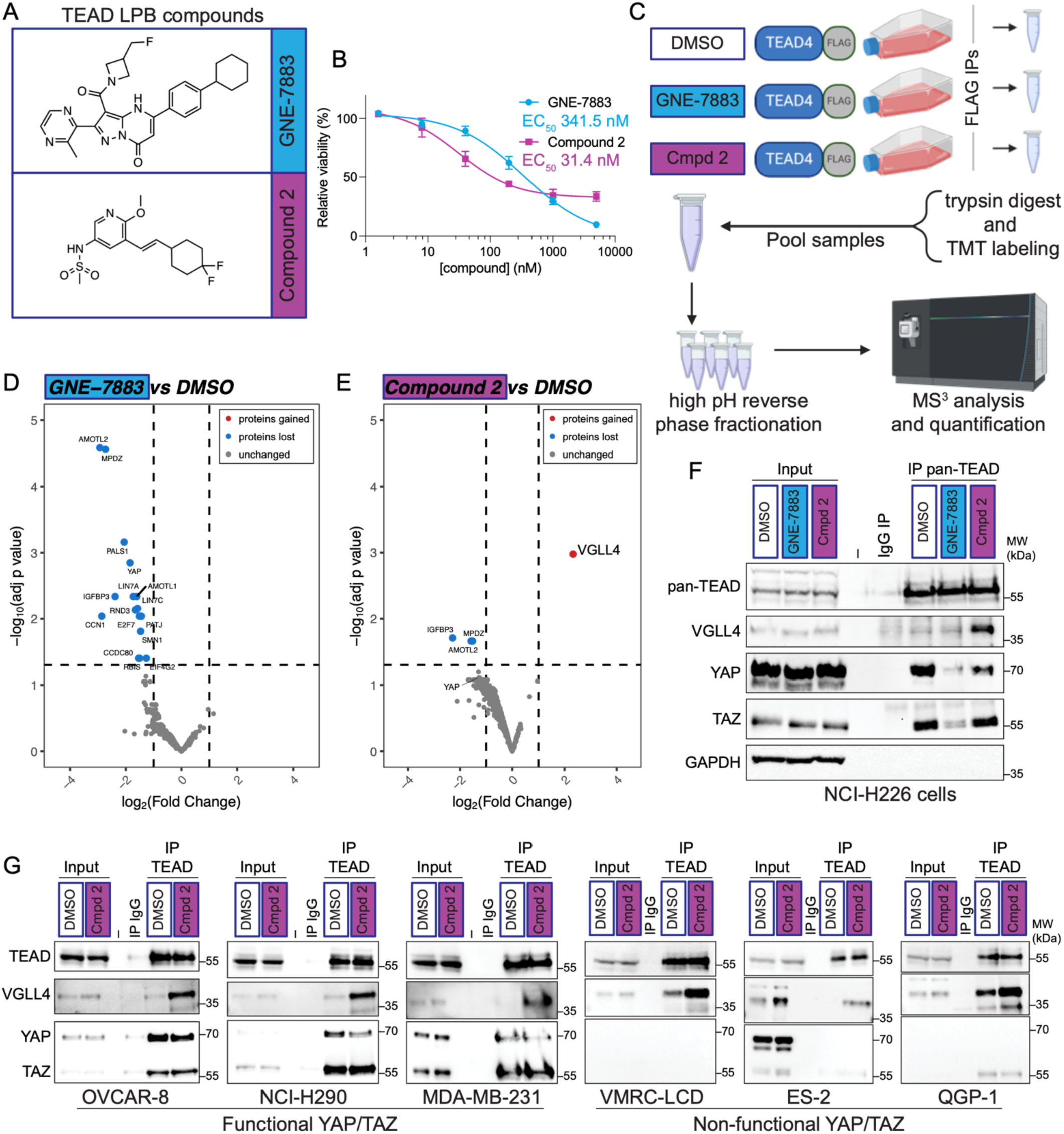
Identification of TEAD interaction partners that are sensitive to LPB TEAD inhibitors. (A) Structures of GNE-7883 and Compound 2. (B) Viability dose response curves (mean ± SD) for GNE-7883 and Compound 2 in NCI-H226 cells treated for six days. n = 3 biological replicates. (C) Schematic of experimental workflow for TEAD-FLAG affinity purification coupled with mass spectrometry (AP-MS) using tandem mass tags (TMT). Three replicates were multiplexed with TMT and fractionated. Mass spectrometry data was collected by LC/MS^3^ using Real Time Search.^79,80^ Data were analyzed using MSstatsTMT.^81^ (D) Volcano plot of TMT proteomic results for TEAD4 AP-MS comparing GNE-7883 treatment to DMSO vehicle control. The TMT-quantified fold-change is plotted against the adjusted *p*-value from model-based testing using MSstatsTMT. Proteins meeting a cutoff of adj *p*-value <0.05 and 2-fold change are highlighted. n = 3 biological replicates. (E) Volcano plot for TEAD4 AP-MS as in (D) but for Compound 2 treatment compared to DMSO. (F) Endogenous pan-TEAD co-immunoprecipitation (coIP) assays in NCI-H226 cells. Cells were treated for 24 h with 3 µM of the indicated compounds or DMSO vehicle control. Cellular extracts were subjected to IP with pan-TEAD antibody or an immunoglobulin G (IgG) control. IP samples were probed for co-precipitating endogenous proteins by immunoblot (IB). Inputs are 10% for pan-TEAD and 1% for others. n = 6 biological replicates. (G) Endogenous pan-TEAD was recovered from lysates of the indicated cell lines treated for 24 h and probed for co-precipitating VGLL4, YAP, and TAZ by IB. Inputs for pan-TEAD are 10% and 1% for others. n = 2 or more biological replicates per cell line. See also Figure S1, Table S1 and Table S2.

To learn how these two small molecules influence TEAD biology we took a quantitative proteomic approach using affinity purification mass spectrometry (AP-MS) centered on TEAD. We used TEAD1 or TEAD4 as baits because these are the TEAD paralogs that are most highly expressed in NCI-H226 cells (Figure S1A). We generated stable cell lines that express inducible, FLAG-tagged forms of TEAD1 or TEAD4, treated the cells with small molecules, recovered proteins by FLAG immunoprecipitation (IP), and took the samples forward for mass spectrometry analysis (Figure 1C and S1B). GNE-7883 treatment reduced the association of 16 proteins with TEAD4, including YAP, AMOTL2, MPDZ, IGFBP3, and CCN1 (Figure 1D). Compound 2 treatment decreased the association of TEAD4 with IGFBP3, AMOTL2, and MPDZ, and increased the association of TEAD4 with VGLL4 (Figure 1E). YAP also decreased with Compound 2 treatment but to an extent that did not reach the significance threshold. Similar AP-MS results were obtained both with TEAD4 (Figure 1D-E, Table S1) and with TEAD1 (Figure S1C-D, Table S2).

The most significantly changed protein in our AP-MS experiments upon treatment with Compound 2 was VGLL4, displaying a five-fold increased association with TEAD4 (Figure S1E) and a seven-fold increased association with TEAD1 (Figure S1F). Increased interaction with VGLL4 was not observed with GNE-7883 treatment. Endogenous coimmunoprecipitations (coIPs) in NCI-H226 cells using a pan-TEAD antibody recovered more VGLL4 when the cells were treated with Compound 2 (Figure 1F). And coIP in six additional cell lines—three with functional YAP and TAZ, and three with nonfunctional YAP and TAZ—also confirmed a boosted interaction between TEAD and VGLL4 upon treatment with Compound 2 (Figure 1G). Thus, interaction between TEAD and VGLL4 can be induced with Compound 2 in a variety of cell types, irrespective of YAP/TAZ presence.

### Sulfonamide-containing compounds promote VGLL4 binding to TEAD

VGLL4 is a direct interacting partner for TEAD that antagonizes YAP/TAZ binding ^22,26,50,51^ but is rarely assayed during the development of TEAD inhibitors. To investigate the molecular determinants that enable Compound 2 to induce the VGLL4–TEAD interaction, we first examined other TEAD LPB molecules. We assayed four published TEAD LPB compounds, MGH-CP1,^52^ VT-107,^41^ TED-347,^53^ and K-975 ^42^ (Figure S2A), and found that none of these compounds induced a change in VGLL4–TEAD association (Figure S2B). We next focused on the chemical backbone of Compound 2 and synthesized four additional compounds that engage the lipid pocket of all four TEADs: N1, N2, S1, and S2 (Methods S1, Figure 2A, Figure S2C). Remarkably, compounds S1 and S2, but not N1 or N2, improve recovery of VGLL4 to levels comparable to that induced by Compound 2 (Figure 2B). A distinct feature common to Compound 2, S1, and S2 is a sulfonamide group appended to the pyridine ring, suggesting the sulfonamide as a critical determinant for promoting VGLL4–TEAD. We next assayed two additional LPB compounds, VT-103 which contains a sulfonamide, and VT-104 which does not contain a sulfonamide (Figure 2C).^41^ Again, sulfonamide-containing VT-103 increased VGLL4 recovery to levels comparable to or higher than Compound 2, while sulfonamide-free VT-104 did not impact VGLL4 recovery (Figure 2D). We also synthesized three additional compounds based on Compound 2 by replacing the sulfonamide group: C2 amino, C2 methyl, and C2 acetyl (Methods S1, Figure S2D). However, replacing the sulfonamide caused a loss in affinity for TEAD proteins (Figure S2E), and we observed no change in VGLL4 recovery by coIP in two Hippo-driven cell lines (Figure S2F-G). Altogether, these results uncover the sulfonamide group as a common feature for the TEAD LPB compounds that promote the VGLL4–TEAD interaction.

**Figure 2:**
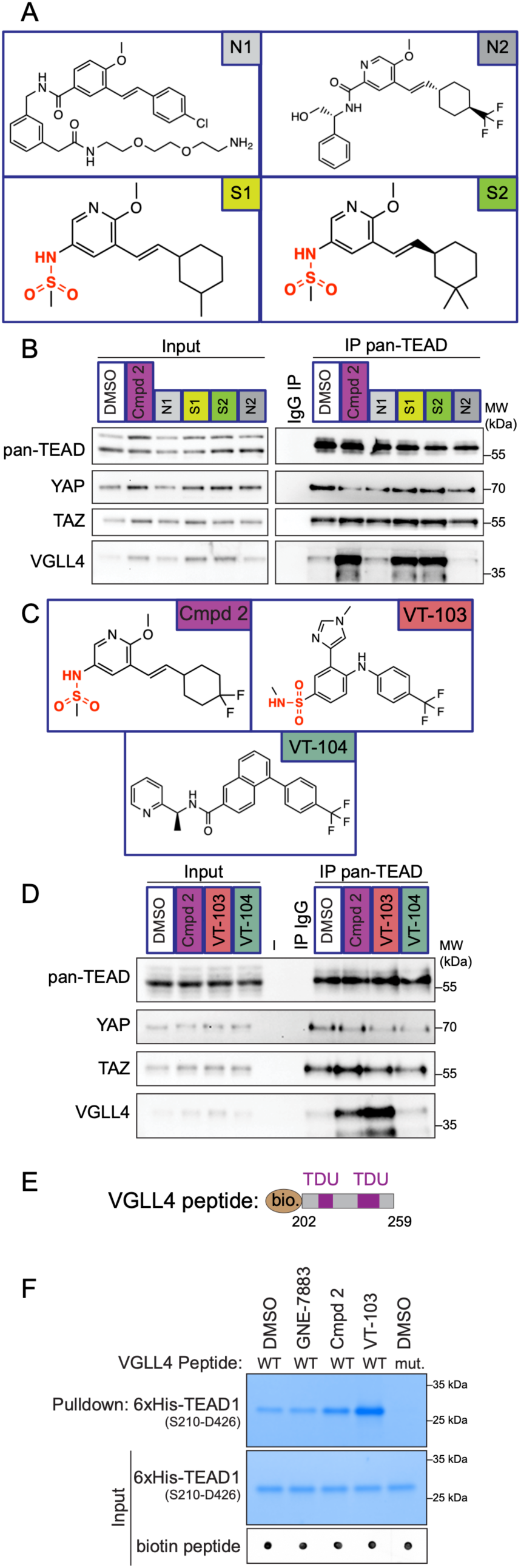
Sulfonamide-containing LPB compounds facilitate the VGLL4–TEAD interaction. (A) Structures of tool compounds without (N1 and N2) and with (S1 and S2) a sulfonamide functional group. Sulfonamide groups are highlighted in red. (B) CoIP experiment to assay TEAD compounds. NCI-H226 cells were treated for 24 h with 3 µM of the indicated compounds or DMSO vehicle control and cellular extracts were subjected to IP with pan-TEAD antibody or an IgG control. Samples were probed with antibodies against the indicated endogenous proteins. Inputs are 10% for pan-TEAD and 1% for others. n = 3 biological replicates. (C) Structures of Compound 2, VT-103, and VT-104. Sulfonamide groups are highlighted in red. (D) CoIP experiments performed as in (B) but treated with the compounds in (C). n = 3 biological replicates. (E) Schematic illustrating the biotinylated (bio.) VGLL4 peptides (residues 202-259) used in the pulldown assays. TEAD interacts with the TDU domains unless these domains are mutated. (F) *In vitro* peptide pulldown assay. Recombinant TEAD1 (S210-D426) was pre-incubated with compounds or DMSO and then incubated with biotinylated VGLL4 peptides, WT or mutant (mut.), bound to streptavidin beads. Recovered TEAD1 was visualized by Coomassie staining. Peptide input was visualized by dot blot. n = 4 biological replicates. See also Figure S2 and Methods S1.

The cocrystal structure of TEAD in complex with Compound 2 shows no notable changes in the overall fold of TEAD that would explain an increase in the VGLL4–TEAD interaction (Figure S2H).^43^ To investigate whether or not a minimal *in vitro* system would reconstitute the gained interaction, we performed a peptide pulldown assay. Using biotinylated VGLL4 peptides (Figure 2E) and recombinant TEAD1 (C-terminal domain), we observed increased recovery of TEAD1 with Compound 2 or VT-103, but not with GNE-7883 (Figure 2F). In fact, VT-103 induced TEAD1 binding to an even greater extent than Compound 2. These *in vitro* findings are consistent with the cellular coIP results from treated cells and indicate that sulfonamide-containing LPB TEAD compounds directly modulate TEAD to induce interaction with VGLL4.

### Compound 2 treatment phenocopies VGLL4 overexpression and counteracts YAP

A key function of VGLL4 is to counteract YAP–TEAD transcriptional complexes to exert growth-inhibitory transcriptional repression.^22,23,27,54^ To determine how cell growth and transcription are affected by enhancing the VGLL4–TEAD interaction, we compared chemical and genetic modulation of VGLL4 using Compound 2 and VGLL4 overexpression respectively. We used Compound 2 because, unlike VT-103 which is TEAD1-specific,^41,55^ Compound 2 is a pan-TEAD molecule. To overexpress VGLL4, we engineered NCI-H226 cells with HA-tagged VGLL4 under control of a doxycycline-inducible promoter, either wildtype or a mutant form of VGLL4 that is unable to interact with TEAD due to mutations made within the two TONDU (TDU) domains (Figure 3A and S3A).^22,27,51^ Doxycycline treatment efficiently induced VGLL4 overexpression (Figure 3B). Simultaneous WT VGLL4 overexpression together with Compound 2 treatment had an additive effect on promoting VGLL4–TEAD association and was accompanied by a concomitant decrease in YAP–TEAD association (Figure S3B), consistent with VGLL4 antagonizing YAP–TEAD complexes. We monitored cellular growth over ten days, and gene expression at 24 hours by probing five representative Hippo target genes: *CTGF* (*CCN2*), *CYR61* (*CCN1*), *ANKRD1*, *F3*, and *IGFBP3*. Both with Compound 2 treatment (Figure 3C) and with WT VGLL4 overexpression (Figure 3D), cellular growth rates slowed and target gene expression decreased. In contrast, overexpression of mutant VGLL4 resulted in little to no change in proliferation or gene expression (Figure 3E). We conclude that Compound 2 treatment is analogous to WT VGLL4 overexpression and that VGLL4–TEAD interaction can promote an anti-proliferative phenotype.

**Figure 3:**
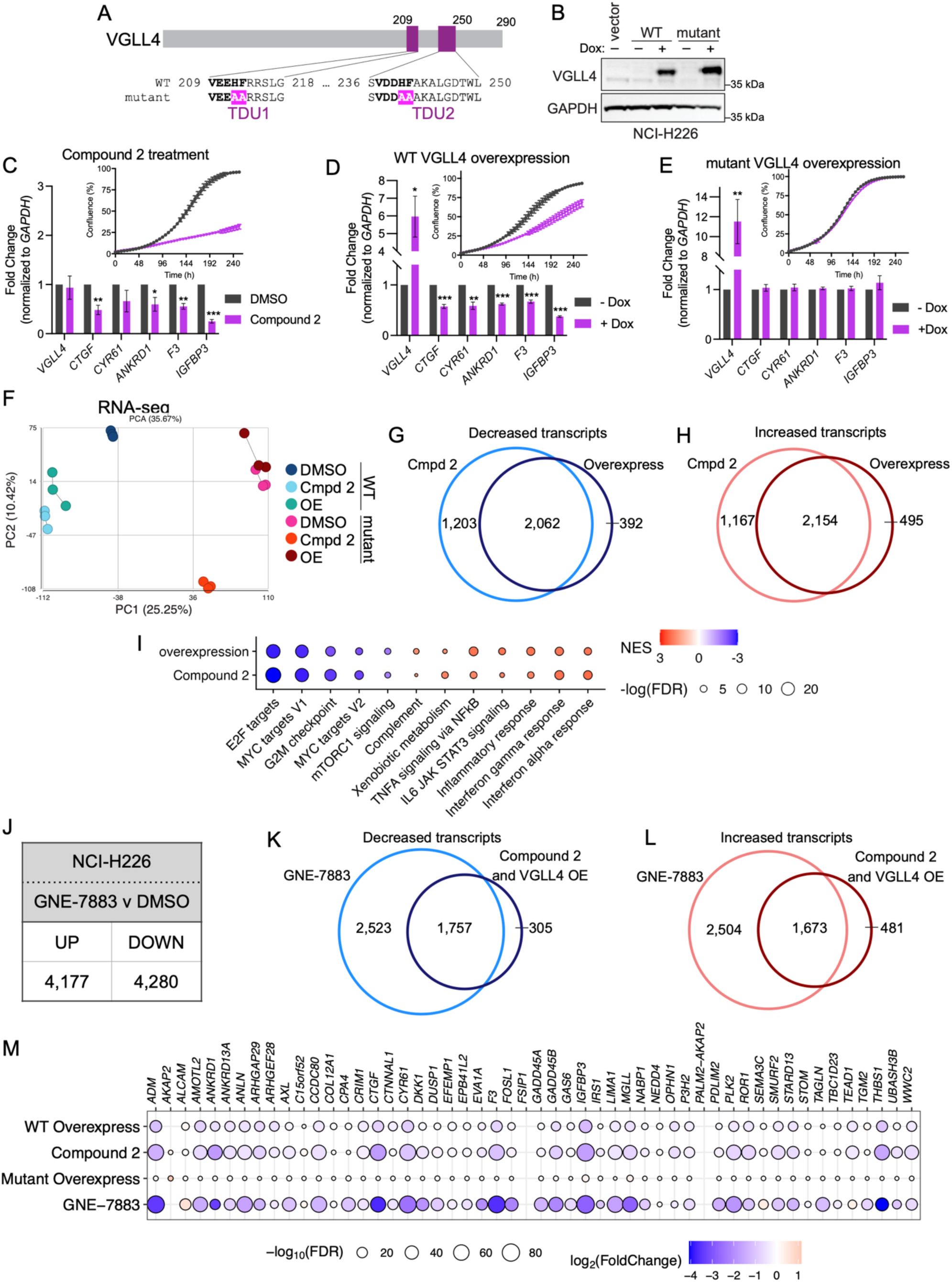
Compound 2 treatment phenocopies VGLL4 overexpression to counteract YAP. (A) Schematic of VGLL4. Amino acid sequences of the two TDU motifs are detailed for WT and the TDU mutant. The four amino acid substitutions that comprise mutant VGLL4 are highlighted in pink (H212A, F213A, H240A, F241A). (B) Immunoblots of doxycycline (dox) induced overexpression of WT and mutant VGLL4 in NCI-H226 cells. (C) Characterization of the impact of Compound 2 treatment on transcription and proliferation in NCI-H226 cells. Cells were cultured in 3 µM Compound 2. For gene expression analysis, cells were assayed after 24 h; n = 3 independent biological replicates, mean ± SEM, \*\*\**p* < 0.001, \*\**p* < 0.01, \**p* < 0.05 by parametric unpaired two-tailed t test. For proliferation, cells were imaged every six hours over ten days; mean ± SEM, n = 3 independent biological replicates. (D) Characterization of the impact of WT VGLL4 overexpression on transcription and proliferation in NCI-H226 cells. Cells were treated with dox to induce overexpression. Data were collected and are represented as in (C). (E) Characterization of the impact of mutant VGLL4 overexpression on transcription and proliferation in NCI-H226 cells. Cells were treated with dox to induce overexpression. Data were collected and are represented as in (C). (F) Principal component analysis (PCA) performed on RNA-seq experiments in WT VGLL4 and mutant VGLL4 cell contexts. Treatment conditions are DMSO vehicle control, Compound 2 (3 µM for 24 h), and VGLL4 overexpression. (G) Diagram showing the overlap between gene expression changes that are decreased at 24 h both with Compound 2 treatment and with WT VGLL4 overexpression. (H) Diagram showing the overlap between gene expression changes that are increased at 24 h both with Compound 2 treatment and with WT VGLL4 overexpression. (I) Enriched Hallmark gene sets ^82^ determined by GSEA of RNA-seq from 24-h Compound 2 treatment or WT VGLL4 overexpression. Twelve top enriched gene sets are shown. (J) Number of transcripts significantly (false discovery rate [FDR] < 0.05) altered by 24 h treatment of cells with GNE-7883 compared with DMSO control. n = 3 biological replicates. (K) Diagram showing the overlap between gene expression changes that are decreased at 24 h both with GNE-7883 treatment and with VGLL4 activity (intersection from 3G). (L) Diagram showing the overlap between gene expression changes that are increased at 24 h both with GNE-7883 treatment and with VGLL4 activity (intersection from 3H). (M) Transcript level changes for 52 Hippo target genes upon the indicated treatments as measured by RNA-seq. FDR is represented by size and fold change is represented by color. See also Figure S3 and Table S3.

To profile the global transcriptional changes that occur with Compound 2 treatment and with VGLL4 overexpression, we performed RNA sequencing (RNA-seq) (Figure S3C). To distinguish changes that depend on interaction with TEAD, we also analyzed the TEAD-interaction deficient mutant VGLL4 cell context. Compound 2 treatment or WT VGLL4 overexpression yielded thousands of significantly changed transcripts, a majority of which were less than two-fold in magnitude (Figure S3D-E). In contrast, overexpression of mutant VGLL4 induced a mere 9 transcripts, one of which is *VGLL4* (Figure S3F). Principal component analysis (PCA) revealed clustering of the WT VGLL4 samples for Compound 2 treatment and for VGLL4 overexpression (Figure 3F), indicating that Compound 2 treatment induced similar transcriptional changes to WT VGLL4 overexpression. Indeed, we observed 84% (2,062 of 2,454) overlap of decreased transcripts (Figure 3G) and 81% (2,154 of 2,649) overlap of increased transcripts in the WT VGLL4 samples (Figure 3H). Gene set enrichment analysis (GSEA) and Reactome pathway analyses further emphasized similarity between WT VGLL4 overexpression and Compound 2 treatment (Figures 3I and S3G-H). For both conditions we observed negative scoring of gene categories related to cellular growth and proliferation (such as E2F targets, MYC targets, and G2M checkpoint, Cell Cycle) and positive scoring for gene categories related to inflammatory signaling (such as Interferon alpha response, Inflammatory response, Immune system).

Finally, we asked how the transcriptional effects we observe upon chemical or genetic induction of VGLL4 compared with inhibition of the YAP–TEAD interaction by GNE-7883 treatment. Treatment of NCI-H226 cells with GNE-7883 resulted in ∼4,000 significantly increased transcripts and ∼4,000 significantly decreased transcripts (false discovery rate [FDR] < 0.05, n = 3) (Figure 3J). Overlapping the VGLL4 consensus transcriptional changes (Figure 3G-H) with changes induced by GNE-7883, we found that ∼80% of VGLL4-influenced changes overlapped with the GNE-7883 data for both increased and decreased transcripts (Figure 3K-L). This result indicates that activating VGLL4 chemically or genetically exerts a similar transcriptional effect to disrupting YAP with GNE-7883. To examine this trend further, we zoomed in on a set of 52 YAP/TAZ and TEAD target genes defined previously ^56^ (Table S3) and observed decreased expression for many Hippo pathway-regulated genes with GNE-7883, Compound 2, and WT VGLL4 overexpression, but not with mutant VGLL4 overexpression (Figure 3M). Overall, these transcriptional trends indicate that YAP–TEAD transcriptional activity can be attenuated by VGLL4, and that a TEAD cofactor switch to VGLL4 can be activated by small molecules like Compound 2.

### TEAD sulfonamide LPB compounds promote VGLL4–TEAD at chromatin

Given that VGLL4 binds to TEAD and influences transcriptional outputs, we next asked how treating cells with sulfonamide compounds alters VGLL4 subcellular localization. We performed biochemical subcellular fractionations in cells treated with TEAD LPB compounds and observed an increase in endogenous VGLL4 protein recovered in the chromatin-associated fraction upon treatment with Compound 2 or VT-103 (Figure 4A). GNE-7883 did not alter VGLL4 localization, but did reduce YAP levels in the chromatin fraction (Figure S4A). Notably, we observed that TEAD is consistently predominantly in the chromatin-bound fraction. We also assayed VGLL4 chromatin localization by immunofluorescence. Pre-treating cells with Triton X-100 detergent prior to fixation permeabilized the cells, removed soluble proteins, and enabled us to image residual chromatin-bound proteins.^57^ Because antibodies to endogenous VGLL4 do not give a specific immunofluorescence signal, we used inducible VGLL4-2xHA expressing cells (Figure S4B). Altogether, treatment with Compound 2 or VT-103 resulted in a significant increase in chromatin-associated VGLL4 (Figure 4B-C).

**Figure 4:**
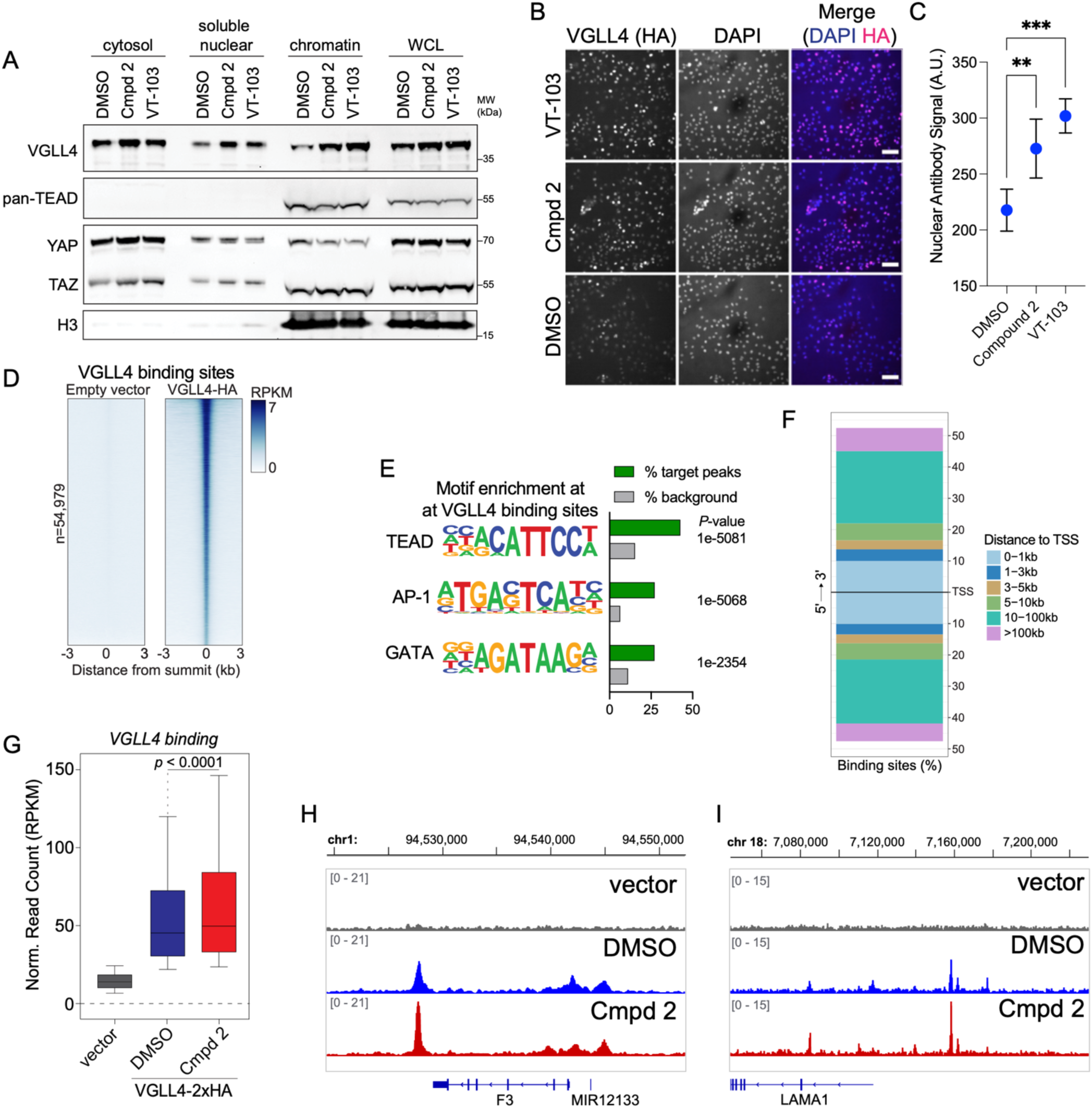
TEAD sulfonamide LPB compounds promote chromatin-bound VGLL4. (A) NCI-H226 cells were treated for 24 h and then fractionated into cytoplasmic, soluble nuclear, and chromatin-associated fractions. Equal amounts of each fraction were analyzed by immunoblotting with the antibodies against the indicated proteins. n = 3 biological replicates. (B) Immunofluorescence staining following triton extraction of soluble proteins. Scale bar is 100 µm. (C) Quantification of triton extraction immunofluorescence experiment in (B). Data are plotted as mean with 95% confidence interval, ordinary one-way ANOVA, ** adj *p*-value = 0.0012, *** adj *p*-value = 0.0008. Note that the y-axis begins at 150. n = 2000 randomly selected cells for each condition. Data are representative of three replicates. (D) Heatmaps for ChIP-seq peaks in cells expressing empty vector control or VGLL4-2xHA. (E) *De novo* motif analysis by HOMER for regions bound by HA-tagged VGLL4. (F) Plot of the peak distribution of the 54,979 VGLL4-HA peaks according to their distance from the nearest annotated transcription start site (TSS). (G) Boxplot showing the ChIP-seq read count signal for the three treatment conditions and assessed by Wilcoxon signed-rank test. n = 3 experiments. (H) Genome browser tracks of VGLL4-HA ChIP-seq at the *F3* locus. (I) Genome browser tracks of VGLL4-HA ChIP-seq upstream of the *LAMA1* locus. See also Figure S4.

To track the genomic binding sites of VGLL4 in NCI-H226 cells, we performed chromatin immunoprecipitation sequencing (ChIP-seq) analysis upon Compound 2 treatment. We monitored VGLL4 using the inducible HA-tagged system, where the levels of epitope-tagged VGLL4 were comparable to endogenous levels of VGLL4 (Figure S4C). We identified 54,979 peaks of VGLL4 binding that were specific to the HA epitope-expressing cells (FDR < 0.01, n=3) (Figure 4D and S4D). Binding sites were enriched primarily for TEAD motifs, as well as AP-1 and GATA sequences (Figure 4E), and the majority of VGLL4 peaks (>70%) were more than 10 kb from the transcription start site of genes, localized in intergenic or intronic regions (Figure 4F and S4E). Upon treatment with Compound 2, we detected enhanced VGLL4 chromatin binding (Wilcoxon Signed-Rank test, p < 0.0001; Figure 4G-I and S4F). Overall, these data indicate that the transcriptional influence of Compound 2 stems from facilitating the VGLL4–TEAD interaction at chromatin.

### VGLL4 overexpression can sensitize cells to sulfonamide LPB inhibitors

We next investigated how VGLL4 protein levels impact the cellular response to TEAD LPB compounds in other cancer cell models. We mined publicly available expression data from DepMap^58^ and found that NCI-H226 cells have higher RNA expression of *VGLL4* than most other cancer cell lines (Figure S5A). We selected five cell types with genetic alterations in the Hippo pathway to cross-compare with NF2-null NCI-H226 cells. NCI-H290 (pleural mesothelioma) and MDA-MB-231 (breast adenocarcinoma) are NF2-null,^8,56^ MSTO-211H (pleural mesothelioma) is LATS-deficient,^59^ OVCAR-8 (ovarian serous adenocarcinoma) is YAP-amplified,^56,60^ and QGP-1 (pancreatic somatostatinoma) is a YAP/TAZ-non-expressing cell line (Figure 1G)^61^ which we selected as a Hippo-independent control. Immunoblotting of lysates showed that VGLL4 protein was notably higher in NCI-H226 cells compared to these other cell types (Figure 5A). We next assayed the sensitivity of these cell lines to TEAD LPB compounds. Although all of the Hippo-dependent cell lines were sensitive to GNE-7883 (Figure S5B), only NCI-H226 cells responded appreciably to Compound 2 and VT-103 (Figure 5B). As expected, the YAP/TAZ-deficient QGP-1 cells were insensitive to all three TEAD compounds (Figure 5B and S5B). The observation that cells with more VGLL4 are more sensitive to Compound 2 and VT-103 further implicates VGLL4 as the determining factor required for mediating an anti-cancer response.

**Figure 5:**
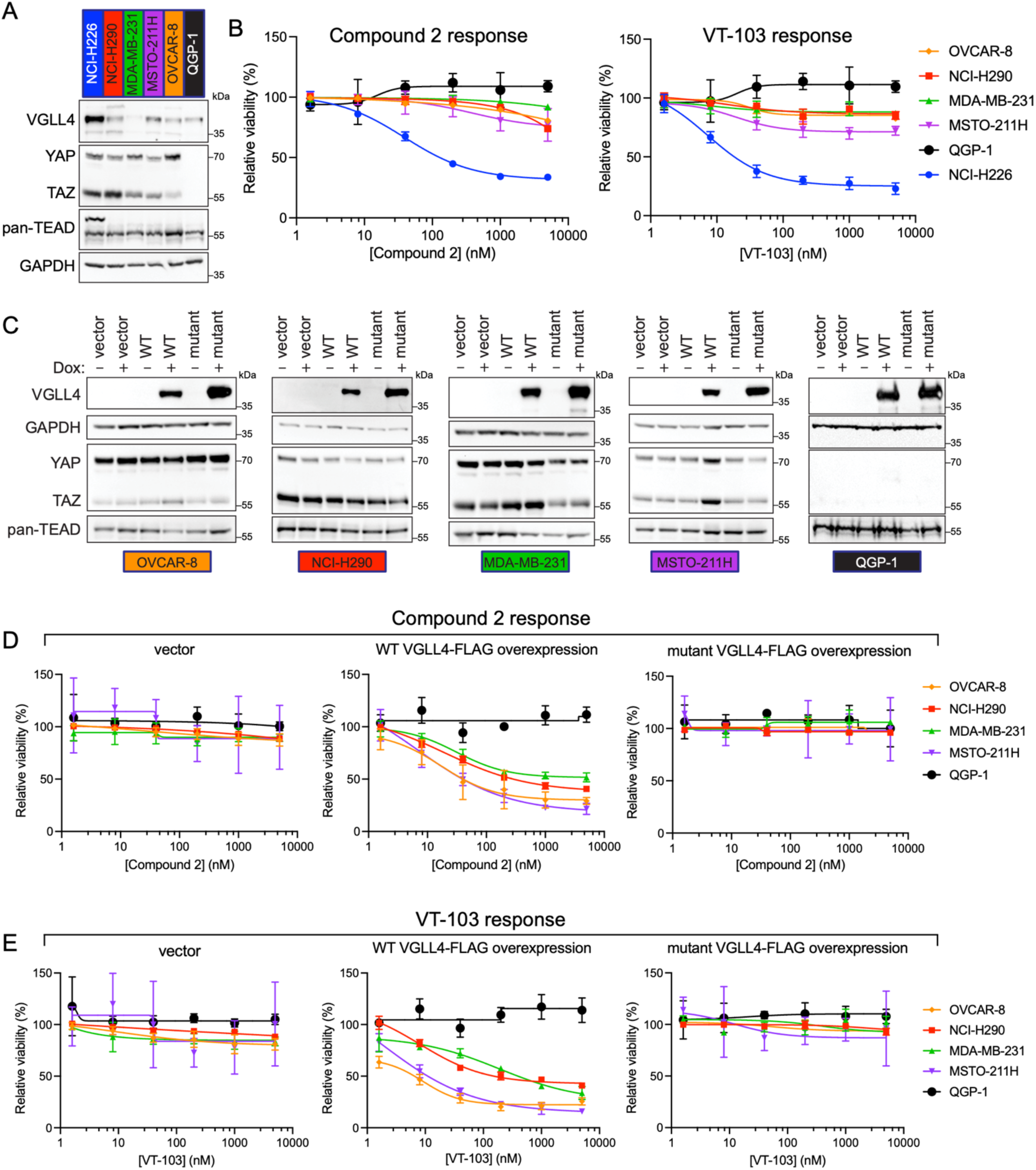
VGLL4 overexpression sensitizes Hippo-dependent cell lines to sulfonamide LPB compounds. (A) Immunoblots of lysates from the six indicated cell lines. (B) Viability dose response curves (mean ± SD) for Compound 2 (left) and VT-103 (right) in the six indicated cell lines treated for six days. n = 3 biological replicates. (C) Immunoblots of lysates from cell lines engineered with inducible VGLL4 overexpression for the five indicated cell types with and without doxycycline (Dox) induction. The VGLL4 mutant used in these experiments is detailed in Figure 3A. (D) Viability dose response curves (mean ± SD) for Compound 2. Viability assays were performed with uniform doxycycline treatment. n = 3 biological replicates. (E) Viability dose response curves (mean ± SD) for VT-103. Viability assays were performed with uniform doxycycline treatment. n = 3 biological replicates. See also Figure S5.

To further test the idea that sensitivity to Compound 2 and VT-103 is linked to VGLL4 protein levels, we next overexpressed VGLL4. We created stable cell lines for six cell types by introducing inducible VGLL4 overexpression constructs, both WT and the TEAD interaction-deficient mutant (Figure 3A, 5C and S5C). When we overexpressed WT VGLL4, the previously-insensitive OVCAR-8, NCI-H290, MDA-MB-231, and MSTO-211H cells showed dose responses to Compound 2 (Figure 5D) and to VT-103 (Figure 5E). With both the vector control and with overexpression of the TDU domain mutant VGLL4, these cell types remained insensitive. In contrast, the Hippo-independent cell type, QGP-1, remained consistently insensitive, even with WT VGLL4 overexpression. We also examined NCI-H226 cells, which were already sensitive to Compound 2 and VT-103, and found that overexpression of WT VGLL4 did not further sensitize cells to Compound 2 or VT-103 (Figure S5D), indicating that endogenous VGLL4 levels exert maximal repressive activity in this cell type. Altogether, these results reveal VGLL4 as the critical factor through which these sulfonamide-containing TEAD-targeting molecules exert anti-cancer efficacy and indicate that YAP activity is necessary to elicit a proliferative response.

### VGLL4 knockout confers resistance to Compound 2 and VT-103 in NCI-H226 cells

Given the antiproliferative effects of inducing the VGLL4–TEAD interaction, we next asked how NCI-H226 cells respond to the opposite perturbation, deletion of VGLL4. CRISPR targeting of an intron-exon junction within the *VGLL4* locus (Figure 6A) resulted in efficient cellular depletion of VGLL4 protein (Figure 6B), with no influence upon cell growth rate (Figure S6A). A pooled transfected population of cells showed similar VGLL4 depletion compared to a clonal population (Figures S6B-C), thus, to avoid clone-specific effects, we performed all subsequent experiments with the VGLL4 knockout (KO) pool. Parental NCI-H226 cells are acutely sensitive to VT-103 and Compound 2 (Figures 6C and S6D). But remarkably, when we knocked out VGLL4, the cells became resistant to these sulfonamide-containing compounds (Figure 6D and S6E) while remaining sensitive to the non-sulfonamide compounds VT-104 and GNE-7883 (Figures 6C-D and S6F). Thus, VGLL4 is necessary for the antiproliferative responses to Compound 2 and VT-103.

**Figure 6:**
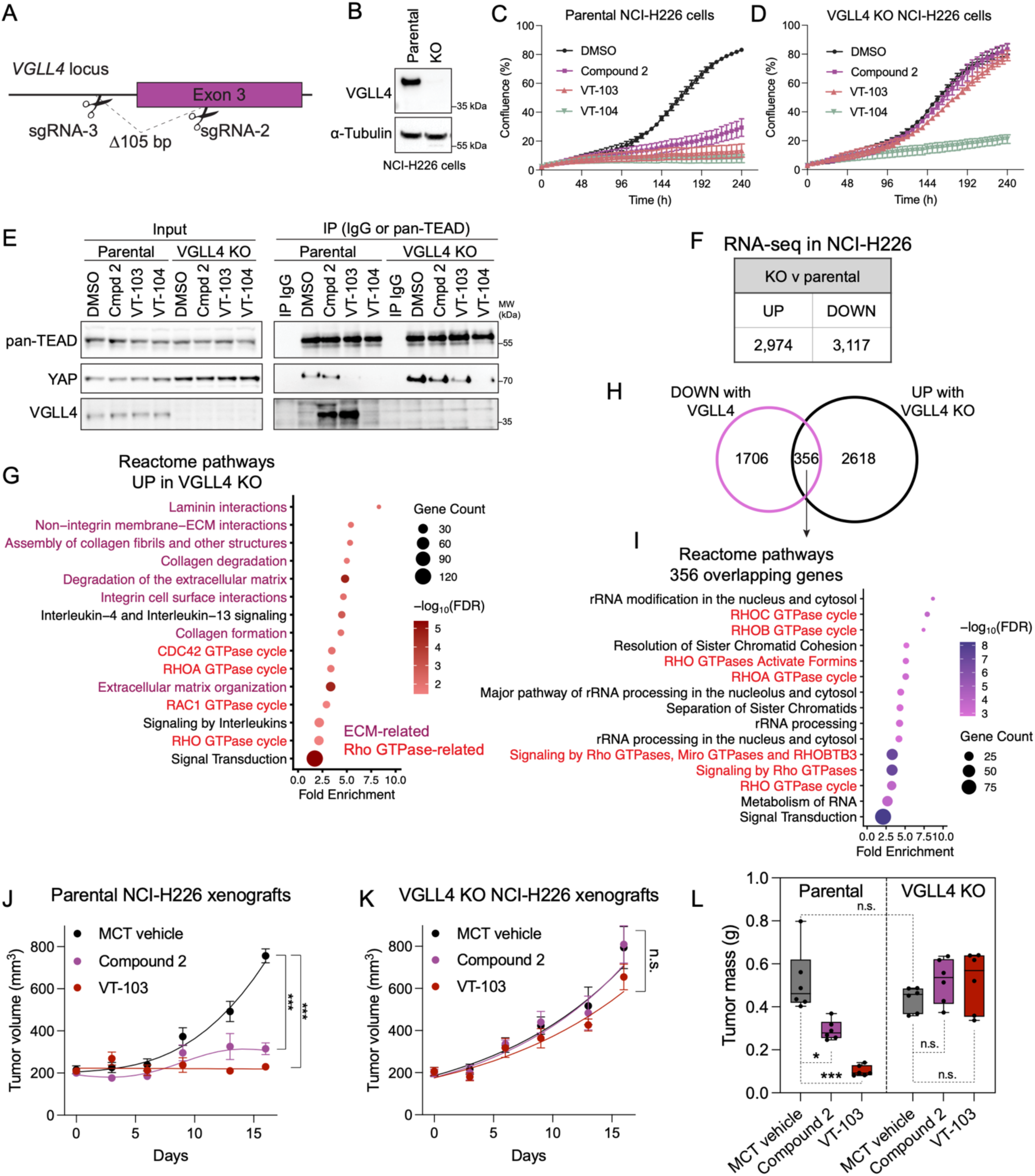
VGLL4 is necessary for the anti-cancer activity of sulfonamide LPB compounds. (A) Schematic of the CRISPR knockout strategy targeting *VGLL4*. (B) Immunoblots of lysates from parental NCI-H226 cells and from a knockout pool of NCI-H226 cells. (C) Proliferation (mean ± SEM) of NCI-H226 parental cells grown in the presence of 3 µM compound or DMSO. n = 3 experiments. (D) Proliferation (mean ± SEM) of NCI-H226 VGLL4 knockout (KO) cells grown in the presence of 3 µM compound or DMSO. n = 3 experiments. (E) Immunoblots of coIP assays from parental and VGLL4 KO cells. Cells were treated for 24 h with 3 µM of the indicated compounds or DMSO vehicle control and cellular extracts were subjected to IP with pan-TEAD antibody or an IgG control. Samples were probed with antibodies against the indicated endogenous proteins. Inputs are 10% for pan-TEAD and 1% for others. n = 3 biological replicates. (F) Number of transcripts significantly (FDR < 0.05) altered by RNA-seq in the VGLL4 KO pool of cells compared to parental NCI-H226 cells. n = 3 biological replicates. (G) Pathway enrichment analysis on the highly significantly increased genes from RNA-seq (FDR < 0.05 and |log_2_FoldChange| > 1). Reactome Knowledgebase ^83–85^ pathways were ranked by FDR and the top 15 pathways are shown. GTPase categories are presented in red text, ECM categories are in purple. Circle color indicates the FDR, size indicates the number of genes. (H) Diagram depicting the overlap of significantly changed transcripts between those increased with VGLL4 knockout and those decreased upon Compound 2 treatment and VGLL4 overexpression (from Figure 3G). (I) Pathway enrichment analysis on the 356 genes from (H). highly significantly increased genes from RNA-seq (FDR < 0.05 and |log_2_FoldChange| > 1). Reactome Knowledgebase ^83–85^ pathways were ranked by FDR and the top 15 significantly enriched pathways are shown. GTPase categories are presented in red text. Circle color indicates the FDR, size indicates the number of genes. (J) Tumor volumes (mean ± SEM) of mice bearing parental NCI-H226 xenograft tumors treated daily with VT-103 at 5 mg/kg, Compound 2 at 200 mg/kg, or MCT vehicle control. n = 6 mice per group. ***p < 0.00001 and by unpaired t tests with Welch correction. (K) As in (J) but for mice bearing KO VGLL4 NCI-H226 xenograft tumors. Endpoint tumor volumes were not significant by unpaired t tests with Welch correction. (L) Tumor masses from mice in (J) and (K). Tumors were excised 16 days after injection and measured. Box plots show spread and averages weights of tumors from each treatment group, n = 6 per group. *p = 0.012, ***p = 0.0001, and ns = not significant, by Welch’s two-tailed t test. See also Figure S6, Table S3, and Table S4.

Because VGLL4 has been shown to antagonize YAP–TEAD interaction and transcriptional activity in a variety of contexts,^21–23,54,62^ we next interrogated YAP in our VGLL4 KO cell system. Not only did we observe that YAP protein levels are higher in the VGLL4 KO cells compared to unperturbed parental NCI-H226 cells (Figure 6E and S6C), we also observed a higher level of the YAP–TEAD interaction in the VGLL4 KO cells compared to the parental cells (Figure 6E). Thus, VGLL4 depletion influences YAP and YAP–TEAD complexes in NCI-H226 cells.

To ask how VGLL4 depletion influences gene expression, we performed RNA-seq in parental NCI-H226 cells and in the VGLL4 KO cells. Overall, we identified ∼6,000 genes that significantly changed in expression (Figure 6F). Of these changes, ∼1,600 were more than two-fold in magnitude (Figure S6G). Because VGLL4 is a transcriptional repressor, we anticipated that many YAP/TAZ/TEAD target genes might be de-repressed by the deletion of VGLL4. However, most of the 52 YAP/TAZ/TEAD target genes ^56^ (Table S3) were only modestly changed in the VGLL4 KO (Figure S6H). Instead, the genes that change significantly are enriched in factors connected to extracellular matrix (ECM) formation and Rho GTPase activity (Figure 6G, colored text). Interestingly, these are pathways connected to mechanosignaling, which is a major regulatory mechanism for YAP activation.^63–66^ We next surmised that genes repressed by VGLL4 would increase in expression when we knock out VGLL4 and decrease in expression when we activate VGLL4 chemically or genetically (data from Figure 3). Intersecting our RNA-seq datasets, we found 356 genes that match these criteria (Figure 6H, Table S4). These 356 genes again show enrichment of Rho GTPase categories (Figure 6I). Taken together, these results suggest that VGLL4 functions to block YAP not only by competing for binding to TEAD, but also by repressing Rho GTPase and ECM genes that activate YAP upstream.

Finally, to test the impact of VGLL4 deletion in a more tumor-relevant system we employed a NCI-H226 xenograft tumor model, which is frequently used to evaluate efficacy of TEAD-targeting molecules *in vivo*.^8,41,42,67,68^ Using parental NCI-H226 cells or VGLL4 KO NCI-H226 cells, we grafted subcutaneous tumors. No difference in the rate of tumor formation was observed between the parental tumors and the KO tumors (Figure S6J). Once tumors were established, we treated mice with vehicle control, Compound 2, or VT-103 for sixteen days. Compared to vehicle control, growth of parental NCI-H226 tumors was inhibited in the Compound 2- and VT-103-treated mice (Figure 6J), but growth of the VGLL4 KO tumors was insensitive to both compound treatments (Figure 6K). In contrast to the responsive parental NCI-H226 tumors, the VGLL4 KO tumors were not smaller between treatment conditions (Figure 6L). All treatments were well tolerated and did not cause body weight loss (Figure S6K). Altogether, these VGLL4 knockout system experiments reveal VGLL4 as the required factor for the efficacy of sulfonamide-containing LPB TEAD inhibitors in slowing cancer growth *in vitro* and *in vivo*.

## DISCUSSION

A predominant strategy for targeting Hippo pathway deregulation in cancer is by developing small molecule inhibitors that disrupt the oncogenic YAP–TEAD interaction. Our data uncover a surprising, alternate mechanism whereby certain TEAD-targeting compounds promote the repressive VGLL4–TEAD interaction. We show that this chemically boosted VGLL4–TEAD interaction induces a cofactor switch from YAP to VGLL4. VGLL4–TEAD complexes exert an anti-cancer effect by dampening YAP activity and modulating key transcriptional networks, including mechanosignaling genes. Our findings provide evidence that TEAD-targeting molecules do not need to block YAP directly to be impactful and can work instead by inducing VGLL4–TEAD complexes. Based on these results, we argue that VGLL4 should be a factor that is evaluated in the development of TEAD-centered therapies.

The data presented here show that sulfonamide-containing TEAD LPB compounds directly influence TEAD to engage VGLL4. Chemically induced VGLL4–TEAD is agnostic of cell type and can be recapitulated *in vitro*. However, the structural mechanism by which these compounds induce VGLL4–TEAD association is currently unclear. One possibility is that the compounds induce an allosteric change that favors a VGLL4-bound state. Displacing the palmitoyl modification with the more rigid LPB compound could alter the conformation of TEAD, or simply reduce the in-solution flexibility of TEAD. Alternatively, the sulfonamide could be acting as a molecular glue to enhance the VGLL4–TEAD interaction. Either way, the compound-induced structural changes are unlikely to be dramatic because published crystal structure data often show no significant changes in TEAD conformation with LPB compounds,^8,44,52,55,69–71^ including for Compound 2.^43^ Indeed, crystal structures often fail to fully explain how LPB molecules disrupt YAP–TEAD,^8,71,72^ and the minimal insights from currently available crystallographic data is a considerable limitation for the field. More interrogation, such as with in-solution structural techniques, is required to understand the impact of LPB compounds at atomic-level resolution and dissect how TEAD cofactor interactions are differentially modulated by different LPB molecules.

A key observation from our study is a mechanosignaling gene signature linked to VGLL4. The VGLL4-regulated genes that we identified in Figure 6 encompass integrins, collagens, fibronectin, and many genes involved in Rho-GTPase signaling. These genes are significant because a rigid extracellular matrix and Rho-GTPase signaling are key mechanical inputs that induce nuclear YAP activity.^24,66,73^ The mechanical influence on YAP activity is separable from LATS signaling, and thus would still be relevant in a Hippo mutant context. VGLL4-mediated repression of these genes would weaken mechanical signals and release this activating input for nuclear YAP. Thus, based on this gene signature, we suggest a dual mechanism for VGLL4 in antagonizing YAP activity: one node directly competes for YAP–TEAD binding at DNA, and the second node transcriptionally restricts the expression of proteins involved in mechanical signals to activate nuclear YAP. In a context where Hippo signaling is influencing the potent capability of YAP to promote cell growth, these two mechanisms together would allow VGLL4 to exert tighter control over (potentially oncogenic) YAP activity. It is well-established that multiple upstream inputs influence YAP activity. Accordingly, downstream regulators are also advantageous, if not imperative, for controlling YAP activity. We propose that VGLL4 is one such downstream safeguard against YAP hyperactivity, acting both directly, by competing for YAP binding to TEAD, and indirectly, by repressing the expression of mechanosignaling genes that promote active nuclear YAP.

Cofactor switching in transcriptional regulation is an established molecular mechanism for integrating different signaling pathways and fine-tuning transcriptional activities.^74–76^ In the data presented here, four lines of evidence support the concept of a chemically induced TEAD cofactor switch from YAP to VGLL4. First, the interaction of VGLL4 and YAP with TEAD is mutually exclusive.^22,27^ Second, treatment with Compound 2 leads to a concomitant decrease in the YAP– TEAD interaction (Figures 1 and 2) which is restored upon depletion of VGLL4 (Figure 6). Third, the transcriptional response to Compound 2 is highly correlated with the transcriptional response to GNE-7883, and we observe a lower magnitude of transcriptional changes with Compound 2 (Figure 3). This is consistent with partial inhibition of YAP–TEAD by Compound 2 (via VGLL4), and a more complete blockade of YAP–TEAD with GNE-7883. Finally, in cell lines with high YAP levels and low expression of VGLL4 (such as MDA-MB-231 and OVCAR-8), inducing exogenous VGLL4 confers sensitivity to Compound 2 and VT-103 (Figure 5), indicating that a certain threshold of VGLL4 protein is required to functionally outcompete YAP from TEAD. These data are consistent with studies demonstrating that VGLL4 and YAP compete for binding to TEAD,^21–23,26,62^ and demonstrate that small molecules can induce VGLL4-mediated transrepression of YAP by cofactor switching.

Considering that small molecules targeting the Hippo pathway are at various stages of pre-clinical and clinical development, one important question raised by this research is how these data inform the therapeutic potential of TEAD-targeting molecules. Our findings present VGLL4 as a relevant determinant for the response to specific sulfonamide-containing TEAD-targeting molecules, but recent research by Kulkarni et al.^77^ suggests that VGLL4 may be a relevant factor for mediating a response to TEAD inhibitors more broadly. They report results of a CRISPR knockout screen where they find that *VGLL4* deletion confers resistance to the non-sulfonamide LPB compound VT-107. In our study, VT-107 treatment did not change the VGLL4–TEAD interaction but did disrupt the YAP/TAZ–TEAD interaction (Figure S2B). These results—combined with the trend that low VGLL4 expression in tumors correlates with poor prognosis ^21,22,29–32^— suggest that VGLL4 may play a larger than expected role in mediating the effects of TEAD inhibitors. But, given the predominantly repressive function of VGLL4, it is also important to consider that if a cancer cell has evolved to inactivate VGLL4 (as frequently occurs), then a therapy that acts through VGLL4 will not work. Loss of VGLL4, acquired or inherent, is a vulnerability. Because of this, perhaps an ideal TEAD-targeting molecule would both increase VGLL4–TEAD and directly disrupt YAP–TEAD, even in the absence of VGLL4. Evaluating the role of VGLL4 within emerging experimental and clinical data from various TEAD inhibitors will better our understanding of VGLL4 in cancer. This could be a critical step towards developing effective therapeutics targeting the Hippo pathway.

### Limitations of the study

Certain limitations are inherent to the experimental designs of this study. Our experiments were performed mostly in cell lines, which do not fully account for the intrinsic variability of human tumors. In our biochemical pulldown experiments, we did not investigate all four TEAD paralogs and instead focused on TEAD1 and TEAD4. Because TEAD paralogs are highly similar to one another,^78^ the interaction networks of different TEADs are likely similar to one another, but we are unable to fully assess this given our focus on TEAD1 and TEAD4. Another limitation is that our AP-MS experiments were likely biased toward more stably bound TEAD interacting proteins, and transient and weakly binding proteins were likely lost in wash steps. Finally, as mentioned earlier, determining the structural basis of the allosteric effects that explain how different TEAD LPB compounds influence cofactor interactions will require further investigations and additional methodologies beyond x-ray crystallography.

## ACKNOWLEDGEMENTS

Graphical abstract and the schematic in Figure 1 were created using BioRender. DepMap data was accessed at https://depmap.org/portal. For assistance with mass spectrometric methods and data collection we thank Tommy K Cheung and Taylur Ma. For advice and discussions we thank Léa Thai-Savard, Ben Walters, Peter Hsu, Benjamin Tombling, Ana Marcu, Laura Keller, and Zhenru Zhou. The ChIP-seq experiments for VGLL4 were carried out by Diagenode ChIP-seq Profiling service. We thank the Genentech cell repository and compound management groups for resource management. We thank Vishva Dixit and the Genentech Postdoc Program for support and training. This work was supported by internal funding.

## AUTHOR CONTRIBUTIONS

Conceptualization, A.D.G.; Methodology, A.D.G., T.J.H., B.D., W.L., G.U., S.P., J.R.Z., J.J.C.; Software, A.D.G., B.D., M.C., S.P., S.V., D.L.; Formal Analysis, A.D.G., B.D., W.L., M.C., B.J.T., S.P., S.V., D.L., J.R.Z.; Investigation, A.D.G., T.J.H., B.D., N.M.K., V.K., S.P.; Resources, T.J.H., B.D., N.M.K., W.L., G.U., J.R.Z., J.J.C.; Data Curation, A.D.G., B.D., W.L., M.C., J.J.C.; Writing – Original Draft, A.D.G. and J.R.L.; Writing – Review & Editing, A.D.G., J.R.L., A.D., T.J.H., J.J.C., M.C., B.D., W.L., G.U.; Visualization, A.D.G., B.D.; Supervision, J.R.L., A.D.; Project Administration, A.D.G.; Funding Acquisition, J.R.L.

## METHODS

### RESOURCE AVAILABILITY

#### Lead Contact

Further information and requests for resources and reagents should be directed to and will be fulfilled by the Lead Contact, Jennie Lill.

#### Materials Availability

All unique and stable reagents generated in this study are available from the Lead Contact with a completed Materials Transfer Agreement.

#### Data and Code Availability

Mass spectrometry proteomics datasets including raw files, metadata, and results tables are deposited at the MassIVE repository (https://massive.ucsd.edu/) under the accession numbers MSV000094636, MSV000094637, and MSV000094638. RNA-seq and ChIP-seq datasets including raw files and metadata are deposited at the NCBI BioProject database (https://www.ncbi.nlm.nih.gov/bioproject/) under the BioProject accession PRJNA1156808. Reviewer access information for these datasets can be found below. This paper does not report original code. Any other information required to analyze the data in this study is available from the lead contact upon request.

APMS TEAD1 : MSV000094637 (PW : tead1apms)

APMS TEAD4 : MSV000094638 (PW : tead4apms)

Reviewer Link for RNA-seq and ChIP-seq : https://dataview.ncbi.nlm.nih.gov/object/PRJNA1156808?reviewer=2tofs9ggvll632j4k1o90qjno6

### EXPERIMENTAL MODEL AND SUBJECT DETAILS

#### Cell Lines

NCI-H226 (RRID:CVCL_1544; male), OVCAR-8 (RRID:CVCL_1629; female), NCI-H290 (RRID:CVCL_A555; male), MDA-MB-231 (RRID:CVCL_0062; female), VMRC-LCD (RRID:CVCL_1787; male), ES-2 (RRID:CVCL_3509; female), QGP-1 (RRID:CVCL_3143; male), MSTO-211H (CVCL_1430; male), were cultured in RPMI with 10% FBS and 10 U/ml Penicillin-Streptomycin (GIBCO 15140122). All cells were cultured at 37°C and 5% CO2 and split every 2-4 days. All cell lines were tested and confirmed negative for mycoplasma.

#### Mice

Animals were maintained in accordance with the Guide for the Care and Use of Laboratory Animals (National Research Council 2011). Genentech is an AAALAC-accredited facility and all animal activities in this research study were conducted under protocols approved by the Genentech Institutional Animal Care and Use Committee (IACUC).

Female C.B-17 SCID (inbred) mice were obtained from Charles River Laboratories at Hollister. All of the mice used in the study were female and 7–10 weeks of age at the start of the study. The mice were fed ad libitum with an autoclaved rodent diet (LabDiet 5010). Mice were housed 3-4 animals per cage in individually ventilated cages within animal rooms maintained on a 14 h/10 h light/dark cycle. Animal rooms were temperature and humidity controlled, between 20.0 and 26.1°C and 30 and 70%, respectively, with 10–15 room air exchanges per hour.

### METHOD DETAILS

#### Plasmids

For piggyBac mammalian expression plasmids, plasmid (pBH) containing a puromycin selection marker were synthesized to contain the desired full-length TEAD or full-length VGLL4 open reading frames downstream of a doxycycline-inducible promoter. For protein expression of TEAD C-terminal domains, plasmids are described previously.^43^

#### Generation of Stable Cell Lines

Stable integration of doxycycline-inducible expression constructs were generated using the PiggyBac system.^86,87^ Plasmids were transfected at a 1:3 ratio of PiggyBac transposon vector:donor vector. NCI-H226 cells were transfected using the Neon transfection system (1230 V, Pulse width: 10 ms, Pulse number: 4). MDA-MB-231 cells, NCI-H290 cells, OVCAR-8 cells, and MSTO-211H were transfected using Lipofectamine 3000. One to three days after transfection, cells were selected for integration events by treating with 1 µg/ml of puromycin. Stable cell lines were cultured in RPMI media containing tet-free FBS and maintained with 1 µg/ml puromycin.

#### Treatment of Cells with Compounds

Unless otherwise indicated, cells were plated in normal media, and the next day media was changed to media containing 3 µM of the indicated compound. Cells were then incubated with compound treatments for 24 h at 37°C with 5% CO2 before being collected for the relevant analysis.

#### Dose Response Viability Assays

Cells were seeded at 1,000 cells per well in Falcon Flat Bottom 96-well plates (Cat# 353219). All outer wells of the plate were filled with PBS to minimize evaporation of cell media. Then 16–24 h after seeding, cells were treated with experimental compounds. Cells were treated with a six-point titration (1:5) of the desired chemical compounds using the HP D300e digital drug dispenser. Cell growth was assessed after six days of treatment using CellTiter-Glo Luminescent Cell Viability Assay (Promega) and the luminescence was read with a EnVision 2104 Multilabel Plate Reader (PerkinElmer). All cell viability data were collected for at least three technical replicates per time point per condition, and at least three independent biological replicates were assayed per experiment. EC_50_ values for the inhibitors were determined by fitting the nonlinear regression curves generated by GraphPad Prism.

#### AP-MS Proteomic Sample Preparation

The experiment was performed for TEAD1 and for TEAD4 using NCI-H226 cells with doxycycline-inducible TEAD1-FLAG or TEAD4-FLAG. Cells were plated in 15-cm dishes with 7×10^6^ cells per plate. After 24 h, media was exchanged to treat cells with 10 ng/ml doxycycline and 3 µM compound or DMSO vehicle control. After 24 h treatment, cells were collected by scraping into cold PBS and pelleted by centrifugation at 180xg for 5 min. Cell pellets were then lysed in Kischkel buffer (50 mM Tris pH 8.0, 150 mM NaCl, 5 mM EDTA, 1% Triton X-100) supplemented with Halt Protease and Phosphatase Inhibitor Cocktail. Whole cell extracts were sonicated for 15 s and then clarified by centrifugation for 10 min. Protein concentrations were measured by Bio-Rad Protein Assay Dye Reagent and normalized to one other. Anti-FLAG M2 magnetic beads (Sigma) were equilibrated in Kischkel buffer and then added to samples, 20 µl bead slurry per sample. Lysates were rotated with M2 beads overnight at 4°C in LoBind tubes (Eppendorf). The next day IPs were washed three times for 5 min with cold Kischkel buffer. Samples were transferred to new LoBind tubes and washed twice with cold PBS. Samples were eluted twice with 100 µl 10% NH_4_OH (RICCA Cat# 631.5-32) by agitation at 1000 rpm for 5 min at 37°C. Samples were analyzed by immunoblotting before being taken forward for sample prep for mass spectrometry.

Samples were dried down by SpeedVac and resuspended in 8M urea/50 mM HEPES pH 8.5. Protein mixtures were reduced in 10 mM DTT (Pierce) for 1 h and alkylated in 20 mM iodoac*e*tamide (Pierce) for 20 min. After alkylation, DTT was added to 20 mM to quench iodoacetamide. Samples were then diluted 4-fold to 2M urea with 50 mM HEPES pH 8.5. Protein digestion was performed by adding 0.5 µg trypsin enzyme (Promega) per protein sample and proteins were digested overnight at 37°C. After digestion, samples were acidified with trifluoroacetic acid and loaded onto C18 stage-tips.^88^ TMTpro labeling was performed on stage-tips. TMTpro labels were reconstituted in anhydrous acetonitrile and diluted 10-fold with 200 mM HEPES pH 8.0. TMT labels were added to samples on stage-tips and centrifuged at 350xg until all liquid had flowed through. TMT flow-through was recovered and passed through a second time. Samples on stage-tips were then washed three times by flowing through 0.1% trifluoroacetic acid. Peptides were eluted twice, first with 50% acetonitrile/0.1% trifluoroacetic acid and second with 50% acetonitrile/20 mM ammonium formate, pH 10. Labeled peptides were then mixed, acidified, and purified by stage-tip. Next, a small-scale MS analysis was performed to confirm that TMT incorporation rate reached >98%. The samples were then fractionated into six fractions using the Pierce High pH Reversed-Phase Peptide Fractionation Kit and dried by SpeedVac.

#### AP-MS Proteomic TMT-based Quantitative Mass Spectrometry

Dried TMT-labeled samples from the AP-MS proteomic experiments were reconstituted in 2% acetonitrile/0.1% formic acid (buffer A). LC–MS analysis was performed on Dionex Ultimate 3000 RSLCnano system (Thermo Fisher Scientific) and an Orbitrap Eclipse mass spectrometer (Thermo Fisher Scientific). Peptides (∼1µg) were resolved on a 25 cm × 75 μm Aurora column packed with 1.6 μm C18 (Ion Opticks) by a 95 min linear gradient of 4% to 30% buffer B (98% acetonitrile/0.1% formic acid) in buffer A (2% acetonitrile/0.1% formic acid) at a flow rate of 300 nL/min. Data were acquired in the synchronous precursor selection (SPS)-MS3 mode together with real time search (RTS).^79,80^

Each duty cycle included an FTMS1 scan in the Orbitrap at 120,000 resolution with a scan range from 350 to 1350 m/z, automatic gain control (AGC) target of 1.0e6 and maximum injection time of 50 ms. MS2 ions were selected using a top speed data dependent mode and fragmented with a CID energy of 35, AGC of 1.5e4, and a maximum injection time of 100 ms. RTS was performed prior to acquisition of MS3 spectra using InSeqAPI software,^89^ which operated similarly to previously published approaches.^79,80^ The following parameters were used for RTS: Uniprot human database April 2021 version, including 42,524 entries of Swissprot sequences of canonical and protein isoforms and of common contaminants; static modifications of cysteine carbamidomethylation (+57.0215 Da) and TMTpro modification on the N termini and lysines (+304.207146); variable modifications of methionine oxidation (+15.9949 Da) and TMTpro modification on tyrosine (+304.207146); trypsin cleavage with allowance of up to 2 missed cleavage events, up to 3 variable modifications per peptide, a precursor ion tolerance of 20 ppm, a fragment ion tolerance of 1.0005 Da with a 0.4 Da offset, TMT SPS-MS3 Mode, and a maximum search time of 100 ms. Protein closeout parameters were enabled at 5 max peptides per protein with a maximum search time of 100 ms; protein closeout was excluded for the following proteins: YAP (P46937), TAZ (Q9GZV5), and VGLL4 (Q14135). Utilizing the Orbitrap Fusion Tribrid’s SPS mode for isolating MS3 ions, the top 8 MS2 precursor ions that met RTS criteria were selected and fragmented via higher collision dissociation energy (HCD) of 55, AGC of 1.5e5, a maximum injection time of 350 ms, isolation width of 1.2 Da, and a resolution of 50,000 at 200 m/z.

Offline MS/MS spectra searches were performed using Comet v.2019.01 with parameters matched to the RTS search. Peptide FDR was filtered to <2% using Linear Discriminator Algorithm. TMTpro reporter ions produced by the TMTpro tags were quantified with Mojave in-house software package by calculating the highest peak within 20 ppm of theoretical reporter mass windows and correcting for isotope purities. MSstatsTMT ^81^ R package was used for statistical analysis as described below.

#### Statistical Analysis of Mass Spectrometry Data

Quantification and differential abundance analysis of AP-MS proteomics data were performed by MSstatsTMT v2.8.0, an open-source R/Bioconductor package. MSstatsTMT was used to create quantification reports and differential abundance analysis reports using the Peptide Spectrum Matches (PSM) as described above. First, PSMs were filtered out if they were (1) from decoy proteins; (2) from peptides with length less than 5; (3) with isolation specificity less than 50%; (4) with reporter ion intensity less than 2^8 noise estimate; (5) from peptides shared by more than one protein; and (6) with summed reporter ion intensity across all channels lower than 10,000. In the case of redundant PSMs (i.e. multiple PSMs in one MS run corresponding to the same peptide ion), only the single PSM with the least missing values or highest isolation specificity of the highest maximal reporter ion intensity was retained for subsequent analysis. Multiple fractions from the same TMT mixture were combined in MSstatsTMT. In particular, if the same peptide ion was identified in multiple fractions, only the single fraction with the highest mean or maximal reporter ion intensity was kept. For AP-MS data, normalization was not performed. The missing intensities were imputed by MSstats v4.8.7.^90^ The normalized reporter ion intensities of all the peptide ions mapped to the protein were summarized into a single protein level intensity in each channel by MSstats. As a final step, the differential abundance analysis between treatments was performed in MSstatsTMT based on a linear mixed effects model per protein. The inference procedure was adjusted by applying an empirical Bayes shrinkage. The model-based test statistics were compared to the Student t-test distribution with the degrees of freedom appropriate for each protein to test the two-sided null hypothesis of no change in abundance. The resulting p values were adjusted to control the FDR using Benjamini-Hochberg’s method.

#### Immunoprecipitation of Endogenous Proteins

Cells were plated to be confluent two days later. If cells were treated, media was changed to media containing the appropriate treatment for the indicated time. Each plate was rinsed twice with cold PBS, and then scraped into Kischkel buffer supplemented with Halt Protease and Phosphatase Inhibitor Cocktail (Thermo Fisher Scientific). Lysates were sonicated for 15 s and cleared by centrifugation for 10 min at 4°C. Protein concentrations were measured by Bio-Rad Protein Assay Dye Reagent. For each IP 3-5 mg of lysate was used as total input. Antibodies used for IPs were 10 µl of pan-TEAD (Cell Signaling Technologies #13295S) or an equivalent amount of Normal Rabbit IgG (Cell Signaling Technologies #2729S). Antibodies and lysates were gently rotated at 4°C overnight, and the next day a 20 µl bed Pierce Protein A/G magnetic beads was added to each sample. HA IPs were performed overnight using HA magnetic beads (Cell Signaling Technologies #11846). IPs were then incubated beads for 2-6 h and then washed four times with 1 mL cold Kischkel buffer, transferring to new tubes before last wash. Samples were eluted with NuPAGELDS sample buffer (Thermo Fisher Scientific) supplemented with β-mercaptoethanol and taken forward for immunoblotting analysis.

#### Immunoblotting Analysis

Samples were heated at 70°C in NuPAGE LDS sample buffer supplemented with β-mercaptoethanol and subjected to electrophoresis on polyacrylamide gels. Proteins were transferred to PVDF membrane and the membranes were blocked in 5% milk. Then membranes were incubated with primary antibodies overnight. The antibodies used are detailed in the Figures and the Key Resources Table. Membranes were washed three times with TBST and incubated with HRP-conjugated secondary antibodies. Immunoblots were developed by ECL with Supersignal West Pico Plus Chemiluminescent Substrate (Thermo Fisher Scientific).

#### Generating Lysates for Immunoblotting

Cells were collected by scraping into PBS. Cell pellets were lysed in RIPA buffer (Thermo Fisher Scientific) supplemented fresh with Halt Protease and Phosphatase Inhibitor Cocktail (Thermo Fisher Scientific). Lysates were incubated on ice for at least 20 min and insoluble material was cleared by 10 min of centrifugation at 4°C. Protein concentrations were measured by Bio-Rad Protein Assay Dye Reagent, normalized, and taken forward for immunoblotting analysis.

#### Chemistry

See Methods S1.

#### Protein Expression and Purification

C-terminal domain domain (CTD) TEAD proteins were purified as previously described with slight modification.^8,36^ TEAD1-CTD (S210-D426), TEAD2-CTD (A217-D447), TEAD3-CTD (Q216-D435) and TEAD4-CTD (Q215-E434) codon-optimized constructs containing TEV Protease cleavable N-terminal His-tags were expressed in *E. coli* BL21 (DE3) cells by autoinduction. Cell pellets were harvested, resuspended in lysis buffer (20 mM Tris pH 8.0, 300 mM NaCl, 10% glycerol, 0.5 mM TCEP, 20 mM imidazole) supplemented with Roche EDTA-free protease inhibitor tablets, and lysed by sonication. Crude lysates were spun at 14,000 rpm for 10 min to remove cell debris, and the cleared lysates were passed over a gravity-packed column with Nickel-Sepharose Fast Flow 6 Resin (Cytivia) that had been pre-equilibrated with lysis buffer. Flow through was reloaded to pass through the column twice total. The column was then washed with 20 column volumes lysis buffer and bound protein was eluted with elution buffer (20 mM Tris pH 8.0, 300 mM NaCl, 10% glycerol, 0.5 mM TCEP, 250 mM imidazole). The eluted proteins were treated with 2.5% w/v hydroxylamine at pH 7.5 for 30 min at room temperature. Proteins were then size-selected using a Superdex 75 120 mL column that had been pre-equilibrated in SEC buffer (20 mM Tris pH 8.0, 100 mM NaCl, 5% glycerol, 5 mM DTT). Fractions containing TEAD protein were pooled and concentrated.

#### TEAD Lipid Pocket TR-FRET Assay

Assay was performed in 384-well plates as described in Hagenbeek et al.^8^ Briefly, N-terminally His-tagged TEAD proteins (YAP-binding domain) were preincubated with compounds for 30 min at room temperature in Assay Buffer (50 mM Hepes pH 7.2, 100 mM NaCl, 0.01% Tween-20, 1 mM DTT). Biotinylated lipid pocket probes were then added to the TEAD–compound mixture and incubated for 60 min at room temperature. Next, a europium-labeled anti-His antibody (Revvity) and XL665-labeled streptavidin (Revvity) were added to the TEAD–compound–probe mixture and incubated for an additional 30 min. TR-FRET values were then measured using a 2105 EnVision Multilabel Plate Reader (Revvity). The potency of compounds was determined using GraphPad Prism by generating an IC_50_ value using a nonlinear four-parameter curve fit. On each plate samples were measured in technical duplicate. The experiment was repeated twice.

#### Peptide Pulldown Experiments

Biotinylated peptides were pre-bound to Pierce Streptavidin Magnetic Beads (20 µl per reaction) by adding an excess of the indicated peptide (5x excess of the binding capacity of the streptavidin beads) and rotating for 2 h at 4°C. TEAD proteins were pre-incubated with compound at a 1:5 molar ratio of TEAD:compound by rotating for 2 h at 4°C. Beads were then washed three times for 5 min with cold Kischkel buffer. Peptide-bound beads were then added to the TEAD mixtures and samples were rotated for 2 h at room temperature. Beads were washed four times for 2 min with cold Kischkel buffer and eluted with LDS sample buffer supplemented with β-mercaptoethanol. Eluted samples were analyzed by SDS-PAGE followed by staining with Colloidal Blue Staining Kit (Thermo Fisher Scientific).

#### Cell Proliferation Assay

To measure cell proliferation 5,000 cells were seeded per well in 24-well plates and treated as indicated. Alternatively, 10,000 cells were seeded per well in 12-well plates and monitored similarly. The cellular confluence in all wells was monitored every 6 h using an IncuCyte ZOOM (Sartorius). In each plate, three technical replicate wells were assayed for each condition. Each well was monitored with 9 images per well. Experiments were performed three times.

#### RNA Extraction and RT-qPCR Analysis

Cells were plated at sub-confluence and the next day media was exchanged to contain the appropriate treatment. After treatment, RNA was purified using the RNeasy Plus Mini kit (QIAGEN) following the manufacturer’s protocol and including the gDNA Eliminator Spin Columns. RNA was reverse transcribed with the High Capacity cDNA Reverse Transcription Kit (Thermo Fisher Scientific) supplemented with RNAse Inhibitor and analyzed by qPCR using gene-specific Taqman probes, TaqMan Fast Advanced Master Mix (Thermo Fisher Scientific), and a QuantStudio 7 Flex Real-Time PCR System (Thermo Fisher Scientific).

#### Preparation of RNA for RNA-Seq

Cells were plated at sub-confluence and the next day media was exchanged to media with the appropriate treatment. RNA was purified using the RNeasy Plus Mini kit (QIAGEN) following the manufacturer’s protocol and using the gDNA Eliminator Spin Columns. RNA was quantified by Nanodrop and submitted to the Genentech NGS Laboratory for library preparation and deep sequencing with 50 bp single end reads.

#### Subcellular Fractionation

Subcellular fractionation was performed similarly to what was described.^91^ A confluent plate of cells was washed twice in PBS, scraped into PBS and pelleted. Cells were resuspended in 250 µL Buffer A (10 mM HEPES, pH 7.9, 10 mM KCl, 1.5 mM MgCl2, 0.34 M sucrose, 10% glycerol, 1 mM DTT, and Halt Protease and Phosphatase Inhibitor Cocktail). 50 µl of the cell suspension was removed as whole cell lysate (WCL). The remaining volume was incubated on ice for 8 min. Samples were then centrifuged at 1,300 x g at 4°C for 5 min. The supernatant (S1 fraction) and pellet (P1 fraction) were separated and S1 was clarified by centrifugation at 14,000xg at 4°C for 10 min. The resulting supernatant (S2 fraction) was collected and the pellet (P2 fraction) was discarded. The P1 fraction was washed once with 500 µL Buffer A and centrifuged 1 minute at 1,300 x g. The P1 fraction was lysed by resuspending in 100 mL Buffer B (3 mM EDTA, 0.2 mM EGTA, and Halt Protease and Phosphatase Inhibitor Cocktail) and incubated for 30 min on ice, followed by centrifugation at 1,700 x g at 4°C for five min. The resulting supernatant (S3 fraction) was separated from the chromatin-enriched pellet (P3 fraction). P3 was washed once with 500 µL Buffer B and resuspended in 400 µl LDS sample buffer. All samples were brought to 400 µl in LDS sample buffer and heated at 70°C for 10 min. Equal volumes of each fraction were taken forward for immunoblotting.

#### Immunofluorescence for Chromatin-Bound Proteins with Triton Extraction

Protocol was based on the method published in Ratnayeke et al.^57^ NCI-H226 cells expressing doxycycline-inducible VGLL4-2xHA were used. Cells were seeded at 12,000 cells per well in 96-well plates that were pre-coated for 4 h with Purecol (Advanced BioMatrix) and with all outer wells of the plate filled with PBS. The next day, media was exchanged to treat cells with 3 µM compounds and with 10 ng/ml doxycycline for 24 h. In order to stain for chromatin-bound proteins, soluble proteins were extracted from cells before fixation. Media was aspirated off and cells were incubated on ice for 1.5 min in ice-cold 0.1% Triton X-100 in PBS, supplemented with Halt Protease Inhibitor Cocktail (Thermo Fisher Scientific), 150 µl per well. Next, cells were fixed by directly adding 50 µl of 16% paraformaldehyde to each well (final 4% paraformaldehyde) and incubating at room temperature for 25 min. Fixative was removed and cells were gently washed three times with PBS. Cells were blocked for 30 min with Blocking Buffer (10% FBS, 1% BSA, 0.1% Triton X-100, 0.01% NaN3 in PBS) and then stained overnight at 4°C with anti-HA-Tag antibody (Cell Signaling C29F4) diluted 1:500 in Blocking Buffer. Primary antibody was removed and cells were gently washed three times with PBS. Cells were then incubated for 30 min with Hoechst 33342 and 1:2000 Donkey anti-Rabbit IgG (H+L) Alexa Fluor 647 secondary antibody diluted in Blocking Buffer. Secondary antibody was then removed and cells were gently washed three times with PBS. Cells were imaged in PBS on a PerkinElmer Opera Phenix using widefield at 10x magnification (binning 2). Images were analyzed in FIJI (ImageJ).^92^

#### Image Analysis

Images were segmented, quantified, and analyzed using custom scripts based on previously published code in MATLAB.^93^ Uneven illumination in each imaged color channel was calculated by averaging the background of many wells, and then corrected via normalization using the calculated profile. A global background value in each field of view was then calculated and subtracted. Nuclei were identified and quantified by calculating the integrated intensity of Hoechst signal; all other fluorescence values were calculated from the median intensity value in each cell. Nuclear masks were segmented by combining both global and adaptive local threshold approaches using the Hoechst channel. Cells near the image borders were removed from analyses, and to increase the fidelity of segmentation, segmented areas that are too large, small, or warped were also removed from analyses. The nuclear mask was then used to quantify intensity levels in the other color channels. Nuclear antibody signal values for 2000 randomly selected cells per treatment condition were analyzed to quantify changes between treatment conditions.

#### Chromatin Immunoprecipitation Sequencing (ChIP-seq)

Cells were plated in 15-cm dishes and the next day media was exchanged to treat cells with 10 ng/ml doxycycline and DMSO or 3 µM Compound 2. After 24 h, media was removed and 15 ml of PBS with 1 mM MgCl_2_ was added; cells were crosslinked with 60 µl of Chip Cross-link Gold Reagent (Diagenode) for 30 min with gentle orbital shaking at room temperature. Fixing solution was removed and cells washed twice with PBS. Next, 20 ml PBS was added to each plate and 37% formaldehyde was diluted to 11% in Fixation Buffer (0.1 M NaCl, 1 mM EDTA, 0.5 mM EGTA, 50 mM HEPES pH 7.6). 2 ml of 11% formaldehyde mixture was added and cells were further crosslinked for 5 min at room temperature. Crosslinking was quenched with 250 mM glycine for 5 min at room temperature. Cells were washed once with cold PBS and cells were scraped into conical tubes. Cells were pelleted at 500 x g for 10 min at 4°C. Cell pellets were stored at −80°C until further processing.

The chromatin was prepared by Diagenode ChIP-seq Profiling service using the iDeal ChIP-seq Kit for Transcription Factors (Diagenode). Chromatin was sheared using Bioruptor sonication device with water cooler (Diagenode) for 20 cycles using a 30 s ON 30 s OFF settings. Shearing was performed in 1.5 ml Bioruptor Pico Microtubes with Caps (Diagenode) with 2 million cells in 100μl. An aliquot of this chromatin was used to assess the size of the DNA fragments obtained by High Sensitivity NGS Fragment Analysis Kit on a Fragment Analyzer (Agilent). For each sample, 10 μg chromatin was immunoprecipitated using 1 µg of anti-HA tag antibody (Abcam). Chromatin corresponding to 1% was set apart as Input. Libraries were prepared manually from input and ChIP’d DNA using MicroPlex Library Preparation Kit v3 /96 rxns (Diagenode) with 96 Unique Dual Indexes for MicroPlex v3 -Set I and Set II (Diagenode). Optimal library amplification was assessed by qPCR and by using High Sensitivity NGS Fragment Analysis Kit on a Fragment AnalyzerTM (Agilent). Libraries were then purified using AMPure XP Reagent (Beckman Coulter) and quantified using QubitTM dsDNA HS Assay Kit (Thermo Fisher Scientific). Finally their fragment size was analyzed by High Sensitivity NGS Fragment Analysis Kit on a Fragment AnalyzerTM (Agilent). Libraries were pooled and sequenced with Illumina technology with paired-end reads of 50 bp length.

#### CRISPR Genome Editing for VGLL4 Knockout Cells

Single guide RNAs (sgRNAs) were designed to disrupt exon 3 of *VGLL4* and cause a 105 bp deletion and frame shifts. The two sgRNA sequences used are CAGCCCCATCGAGCGCGCTG and TTCAGGGGGCGTTTTCTCAA and were used together at a 1:1 ratio. Chemically modified sgRNAs were synthesized (Synthego) and CRISPR reagents and targeting vectors were delivered to cells using the Neon Electroporation Transfection System (Thermo Fisher Scientific). Reactions of Cas9-sgRNA ribonucleoprotein (RNP) complexes were formed four at a time by combining recombinant Cas9 (Thermo Fisher Scientific) and sgRNAs at a 1:3 molar ratio in Neon buffer R (Invitrogen) and incubating at room temperature for 10 min. RNP complexes were stored at 4°C until use. Electroporation reactions were performed with 300,000 NCI-H226 cells, 2.5 pmol Cas9, and 7.5 pmol sgRNAs. Electroporation reactions were performed using 10 µl Neon tips with the conditions of 1230 V, 10 ms pulse width, and 4 pulses. Cells were immediately placed into warm, antibiotic-free RPMI supplemented with FBS and allowed to recover for two days. Cells were expanded and a subpopulation was sorted for single cell clones using a BD LSR Fortessa (BD Biosciences-US). Populations and sorted clones were analyzed by immunoblotting and qPCR to monitor *VGLL4* expression levels.

#### Mouse Xenograft

NCI-H226 cells, parental or VGLL4 KO, were cultured *in vitro*, harvested in log-phase growth, and resuspended in 1:1 ratio by volume of Hank’s Balanced Salt Solution (HBSS) and Matrigel (BD Biosciences) for *in vivo* inoculation. Female C.B-17 SCID mice were subcutaneously inoculated with 10 × 10^6^ NCI-H226 cells in the right flank. Half of the mice received parental NCI-H226 cells, and half received VGLL4 KO NCI-H226 cells. Tumors were allowed to grow to a volume in an initial range of 150 to 300 mm^3^ before mice were randomized to treatment groups at the start of dosing to create closely matched baseline average tumor sizes across groups. Mice were dosed by oral administration (PO) with 200 mg/kg Compound 2, 5 mg/kg VT-103, or MCT vehicle control once daily. Tumor sizes and body weights were recorded twice weekly over the course of the study. Mice with tumor volumes ≥2000 mm^3^ or recorded body weight loss ≥20% were promptly euthanized. Tumor volumes were determined using digital calipers (Fred V. Fowler Company, Inc, Newton, MA). Tumor volumes were calculated as: tumor size (mm^3^) = 0.5 × longer measurement × (shorter measurement^2^). Percentage animal weight changes were calculated as: body weight change (%) = [(current body weight/initial body weight) − 1) × 100]. Analyses of the growth of tumors were performed using a package of customized functions in R (version 3.6.2; R Foundation for Statistical Computing), which integrates software from open-source packages as described.^94^

### QUANTIFICATION AND STATISTICAL ANALYSIS

Statistical comparisons between replicates for RT-qPCR and image quantification were performed with GraphPad Prism software 10. The n indicates number of biological replicates or individual number of cells for microscopy. The n, error bar representations, and details of statistical tests can be found in the figure legends or under the specific Methods heading.

#### Proteomic Data Analysis

This information can be found under the appropriate methods heading.

#### Ontology and Categorization

Gene ontology enrichment was performed using the resources at https://geneontology.org/. Gene ontology (GO) enrichment analysis was performed with PANTHER Overrepresentation Test (Released 20240226).^95–97^ Reactome Pathway Enrichment analyses were performed using Reactome version 86 Released 2023-09-07 and the significantly enriched categories were ranked by FDR unless otherwise specified. Plots of enrichment analyses were generated using R.

#### RNA-seq Data Analysis

RNA-sequencing data were analyzed using HTSeqGenie ^98^ in BioConductor ^99^ as follows: first, reads with low nucleotide qualities (70% of bases with quality <23) or matches to rRNA and adapter sequences were removed. The remaining reads were aligned to the human reference genome (GRCh38.p10) using GSNAP ^100,101^ version ‘2013-10-10-v2’, allowing maximum of two mismatches per 75 base sequence (parameters: ‘-M 2 -n 10 -B 2 -i 1 -N 1 -w 200000 -E 1 -- pairmax-rna=200000 --clip-overlap’). Transcript annotation was based on the Gencode genes data base (human: GENCODE 27). To quantify gene expression levels, the number of reads mapping unambiguously to the exons of each gene was calculated.

Differential gene expression analyses were performed off of the raw counts matrix after trimmed mean of M-values (TMM) normalization ^102^ and filtering out of low abundance genes. The voom function of the limma package ^103,104^ was implemented to test for significant differences in means of count data. We performed quality weighting and robust empirical Bayes shrinkage to protect against outlier samples or genes. The significantly changed genes were chosen with the criteria FDR < 0.05.

#### ChIP-seq Data Analysis

ChIP-seq results were analyzed using the ENCODE ChIP-seq pipeline (v2.2.1).^105^ ChIP-seq reads were aligned to the human reference genome (hg38) using Bowtie2 (v2.3.4.3).^106^ Aligned reads were then filtered for quality and duplicates using samtools (v1.9) ^107^ and Picard (Broad Institute - v2.20.7). The SPP peak caller was used to call ChIP-seq peaks for VGLL4, and input was used to assess the background of the experiments.^108^ Peak sets were filtered using a list of genomic regions that contain anomalous, unstructured, or experiment independent high signal.^109^ ChIP-seq bam and bed files were then used to call differential peaks in the genome by using DiffBind (v3.12.0).^110^ Briefly, Differential Binding Analysis object was created by loading the bam and bed files with the dba() function. Reads were counted with the dba.count() function, followed by depth normalization using the dba.normalize() function, and differential peaks were called across the conditions with the dba.analyze() function using DESeq2 and the following statistical parameters (FDR<0.01). Differential VGLL4 peaks were annotated to the nearest genes by ChIPseeker (v1.38.0),^111^ and de novo transcription factor motif analysis was performed with HOMER.^112^ Read distribution was plotted using deepTools2,^113^ using a 6 kilobases long genomic window that is centered on the identified peaks’ summits, and read enrichment was quantified on a boxplot for which, read coverage was computed by deepTools2’s multiBamSummary function, and statistically significant changes were determined by Wilcoxon Signed-rank tests. Tracks were visualized using IGV (v2.17.1).^114^

## KEY RESOURCES TABLE

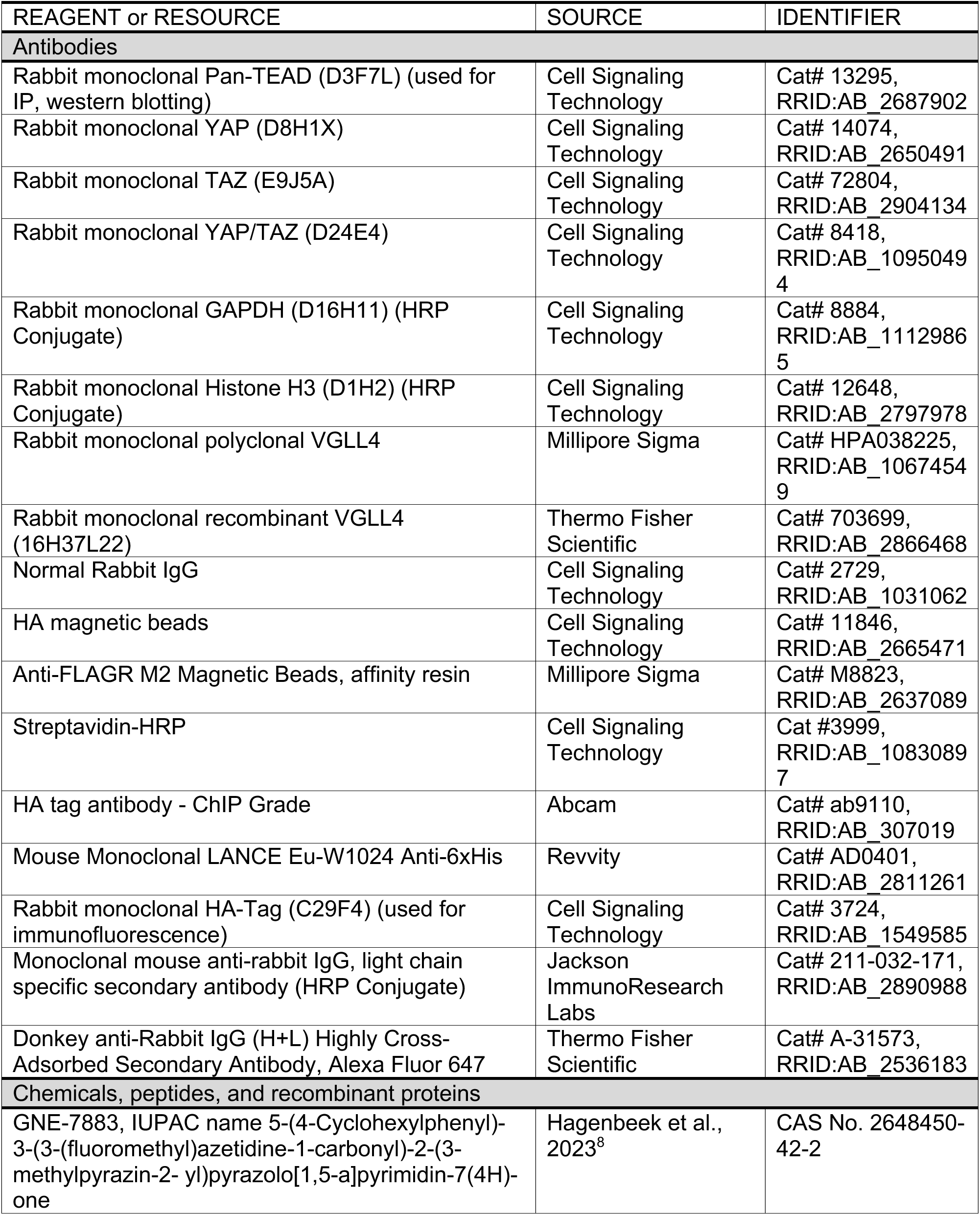
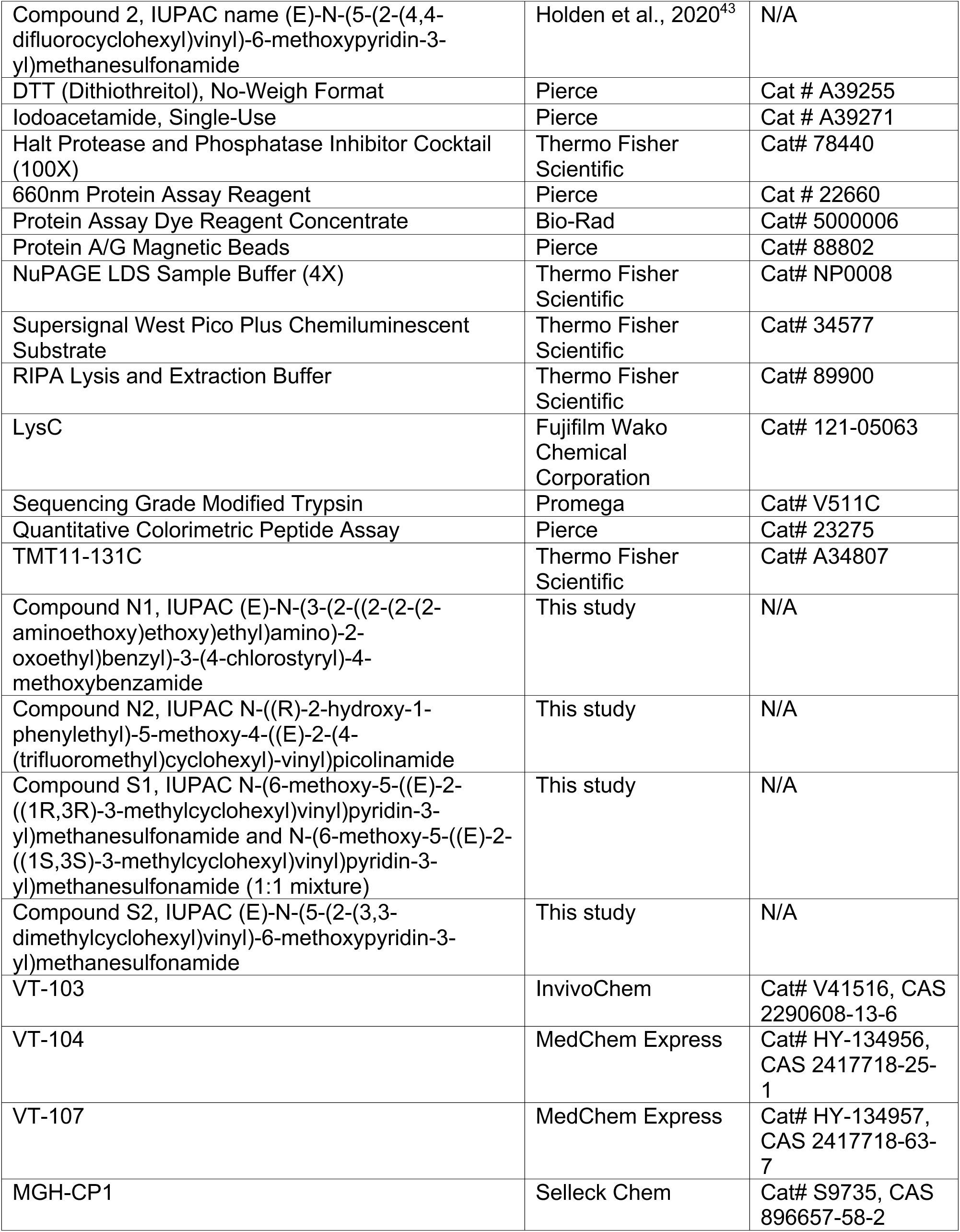
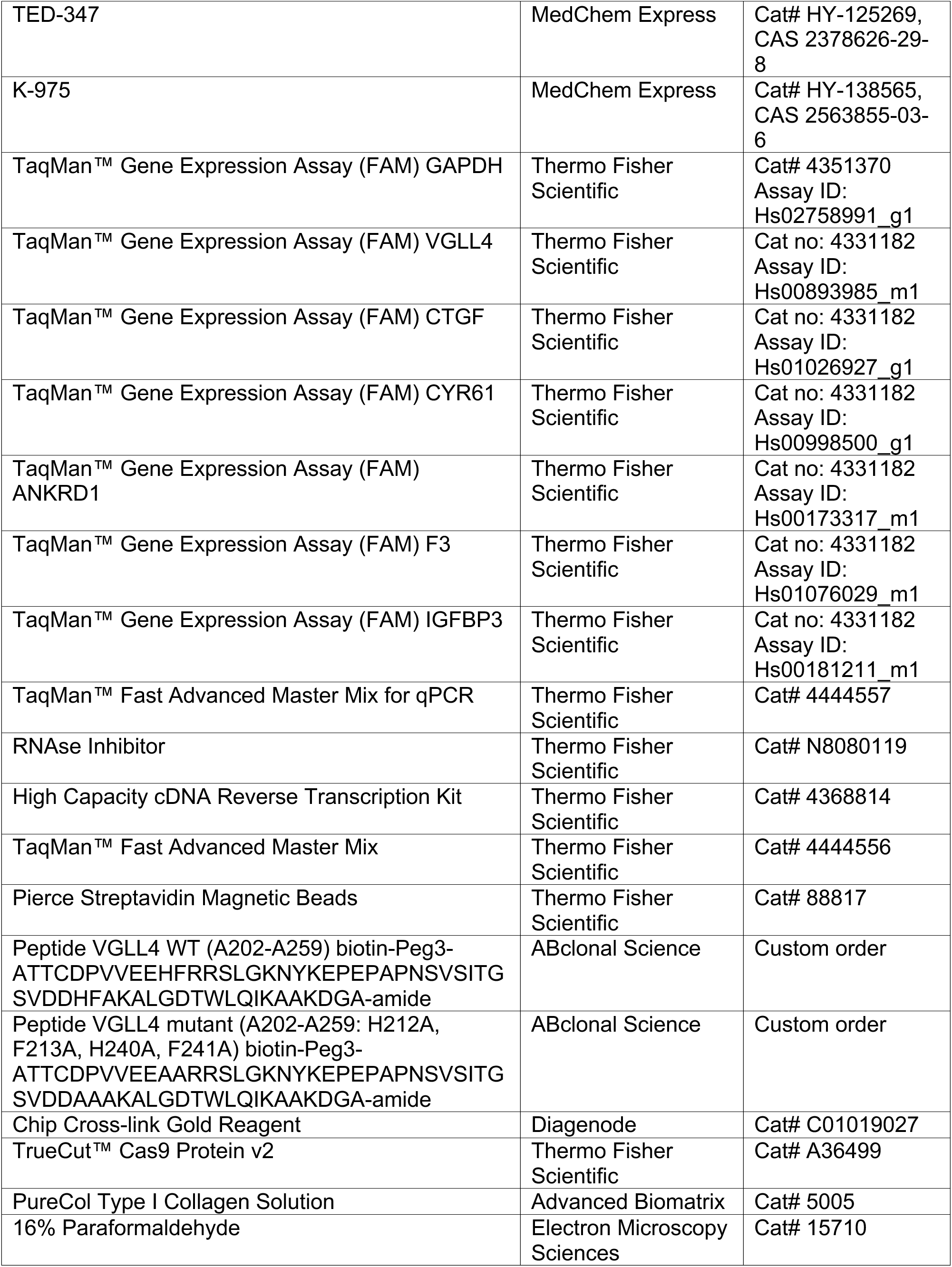
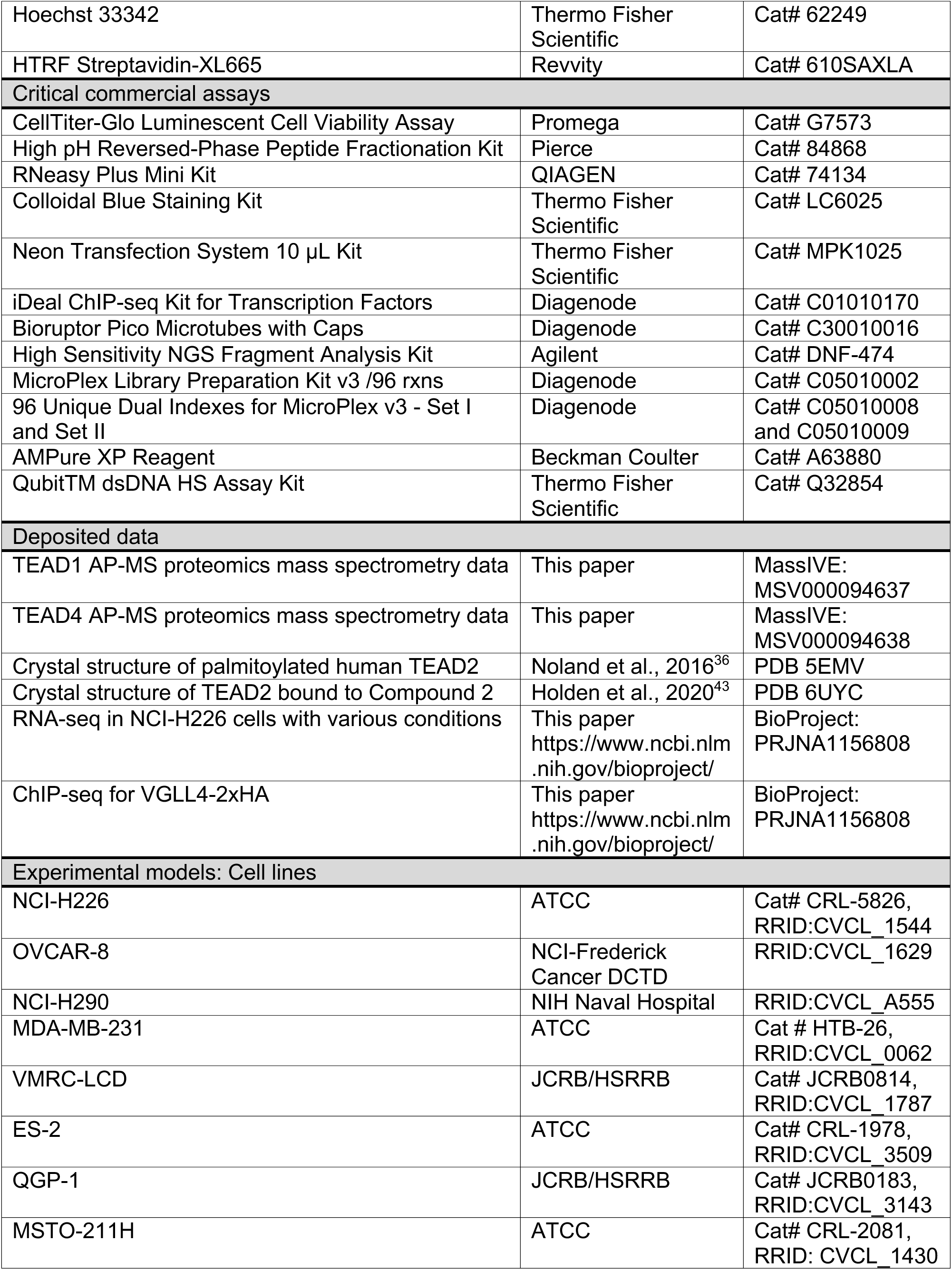
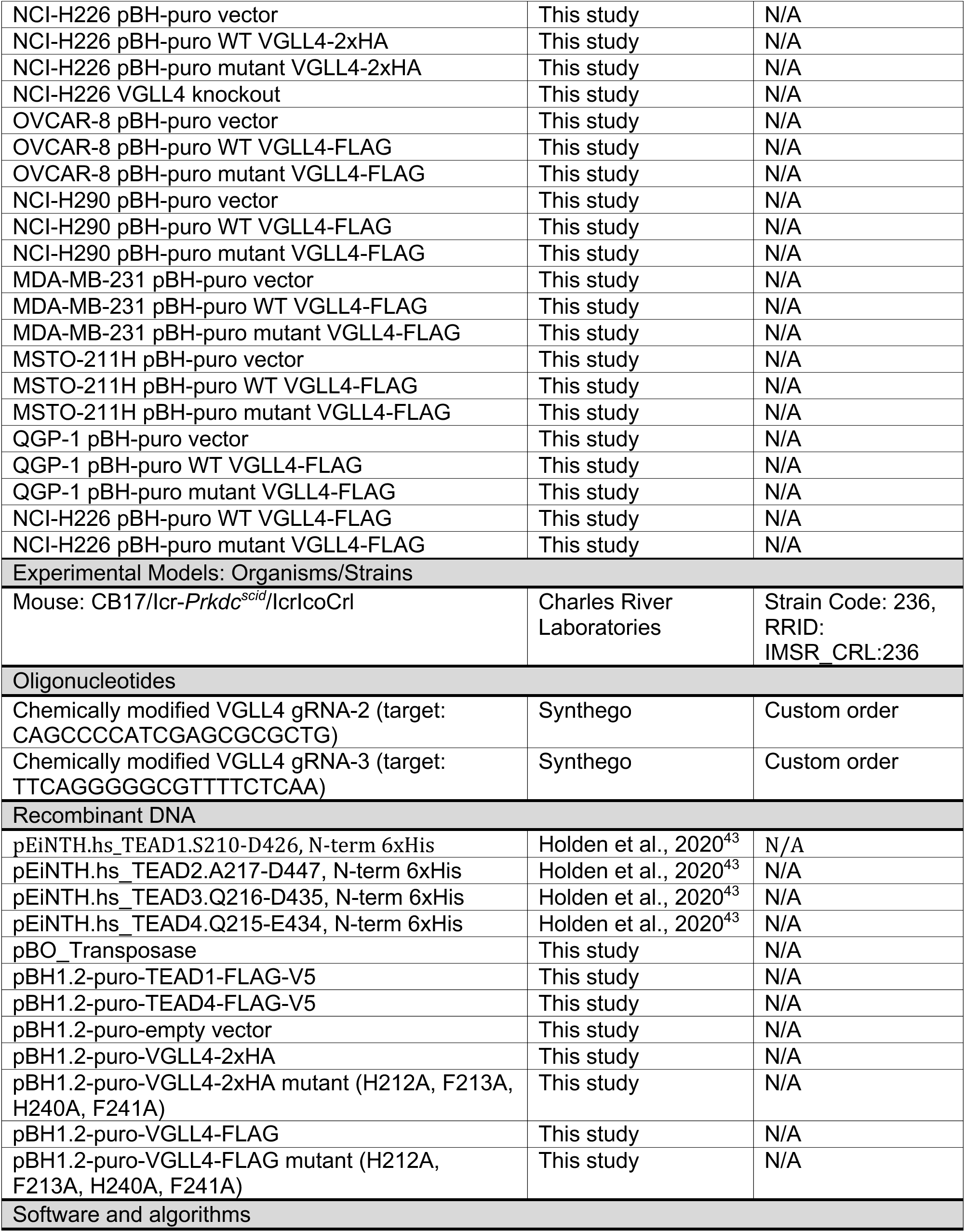
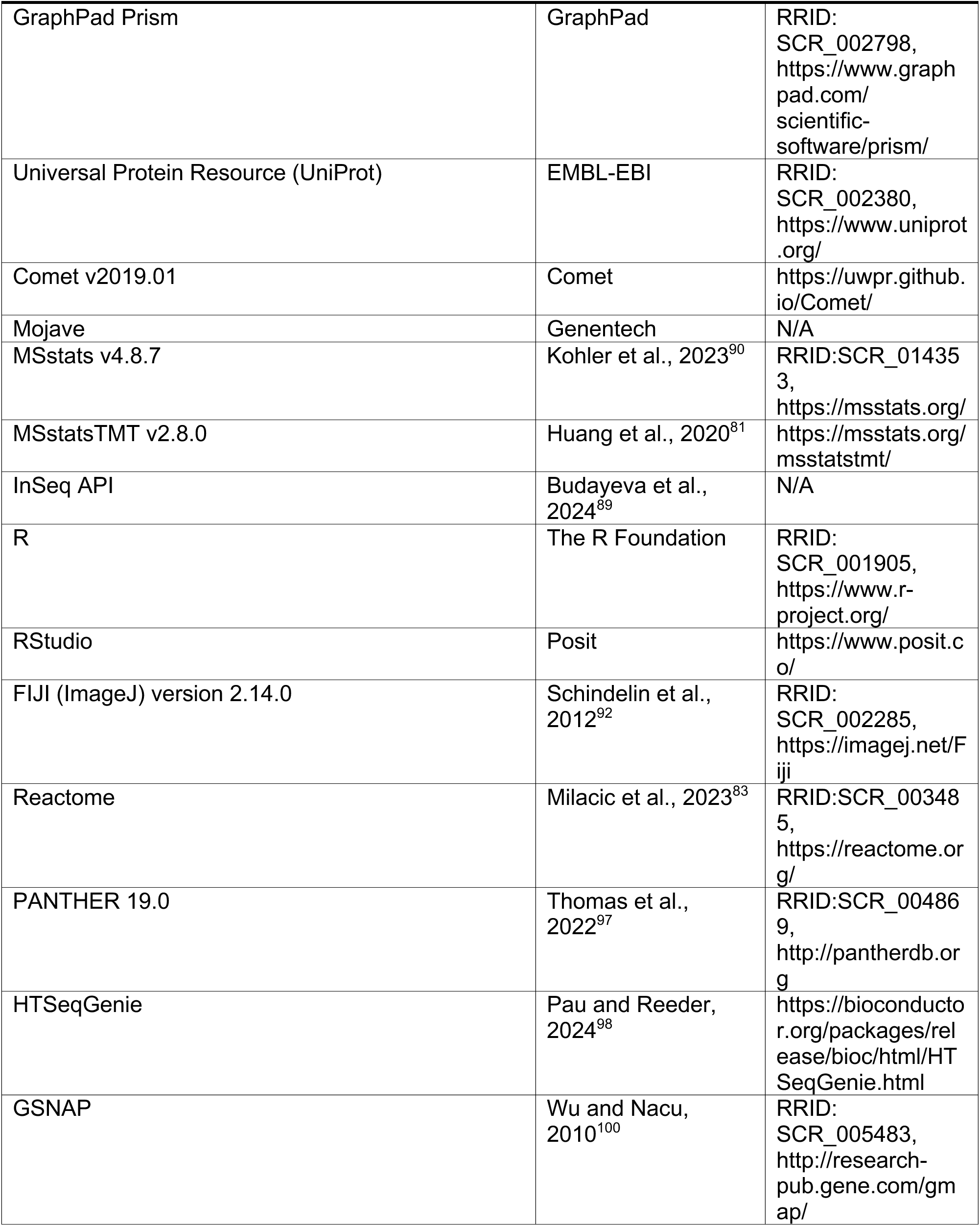
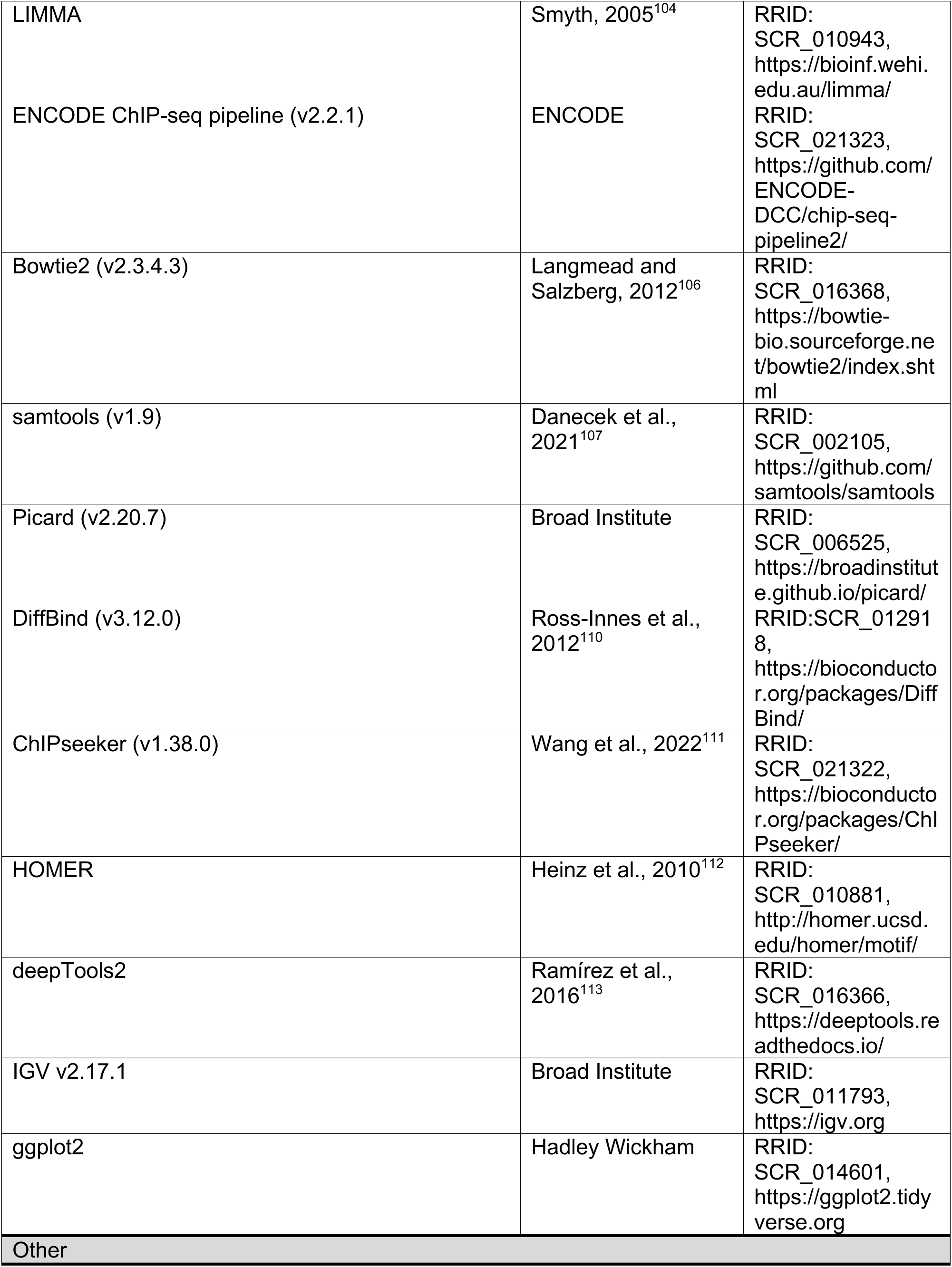
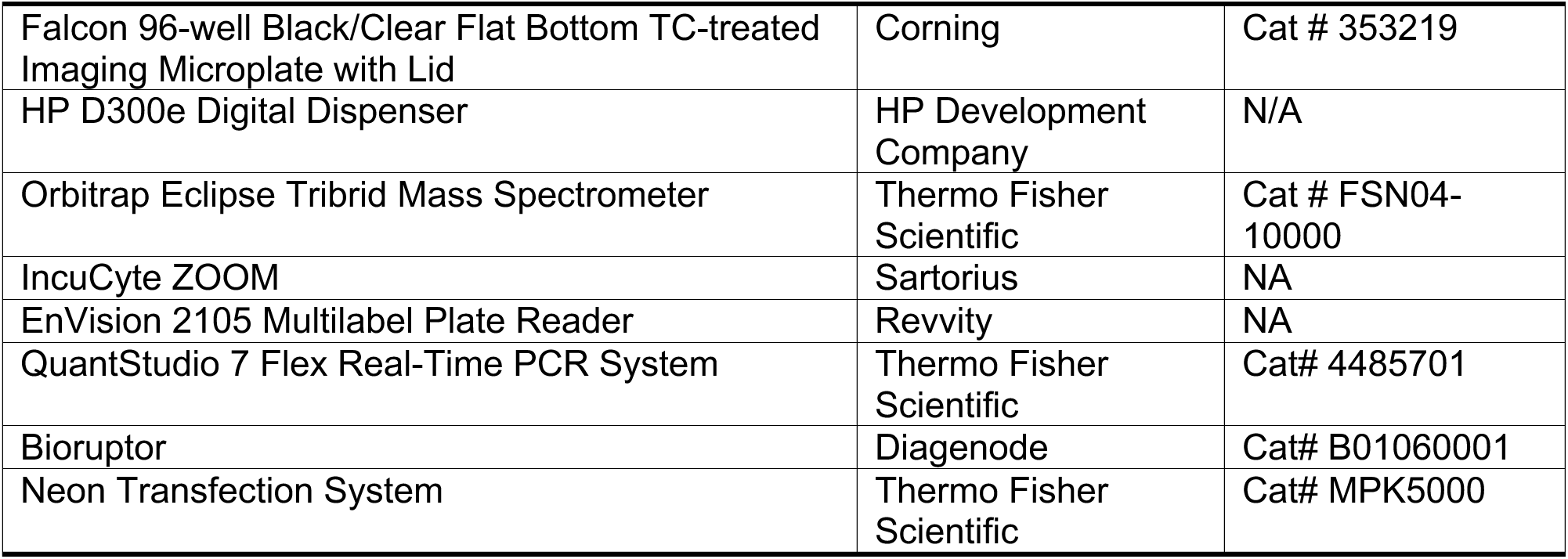

## SUPPLEMENTAL INFORMATION

**Figure S1:**
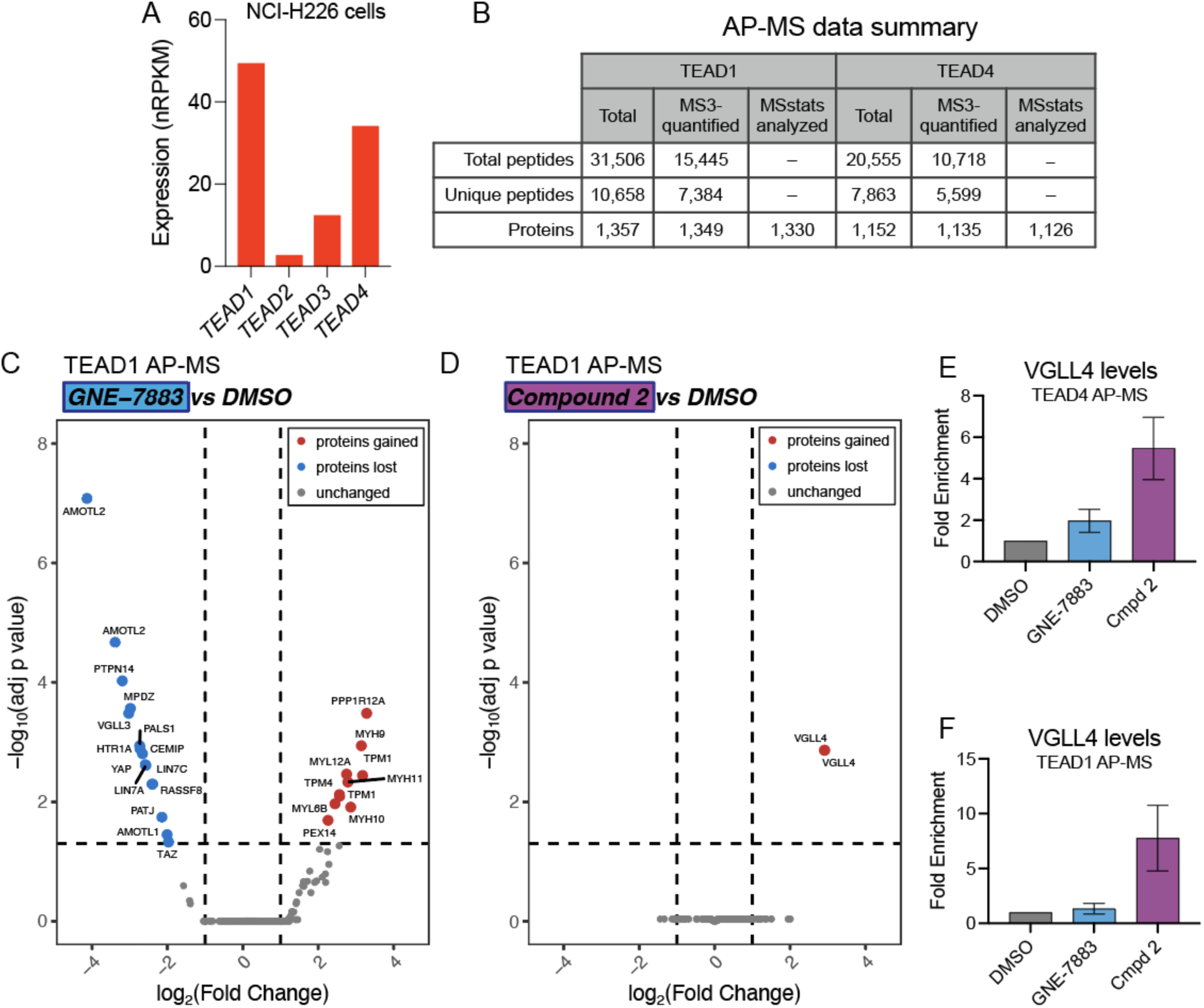
Proteomic analysis of TEAD inhibition. Related to Figure 1. (A) Graph of RNA-seq expression values for all four TEAD paralogs in NCI-H226 cells. (B) Table describing the summary of peptides and proteins analyzed by AP-MS for the TEAD1 experiment and the TEAD4 experiment. (C) Volcano plot of TMT proteomic results for TEAD1 AP-MS comparing GNE-7883 treatment to DMSO. The TMT-quantified fold-change is plotted against the adjusted *p*-value from model-based testing using MSstats. Proteins meeting a cutoff of adj *p*-value <0.05 and 2-fold change are highlighted. n = 3 biological replicates. (D) Volcano plot for TEAD1 AP-MS as in (C) but for Compound 2 treatment compared to DMSO. (E) Fold enrichment relative to DMSO for the TMT-quantified levels of VGLL4 protein (UniProt Q14135) co-purified with TEAD4 in the AP-MS experiment. Data are represented as mean ± SD; n = 3 biological replicates. (F) As in (E) but for the TEAD1-affinity purified samples.

**Figure S2:**
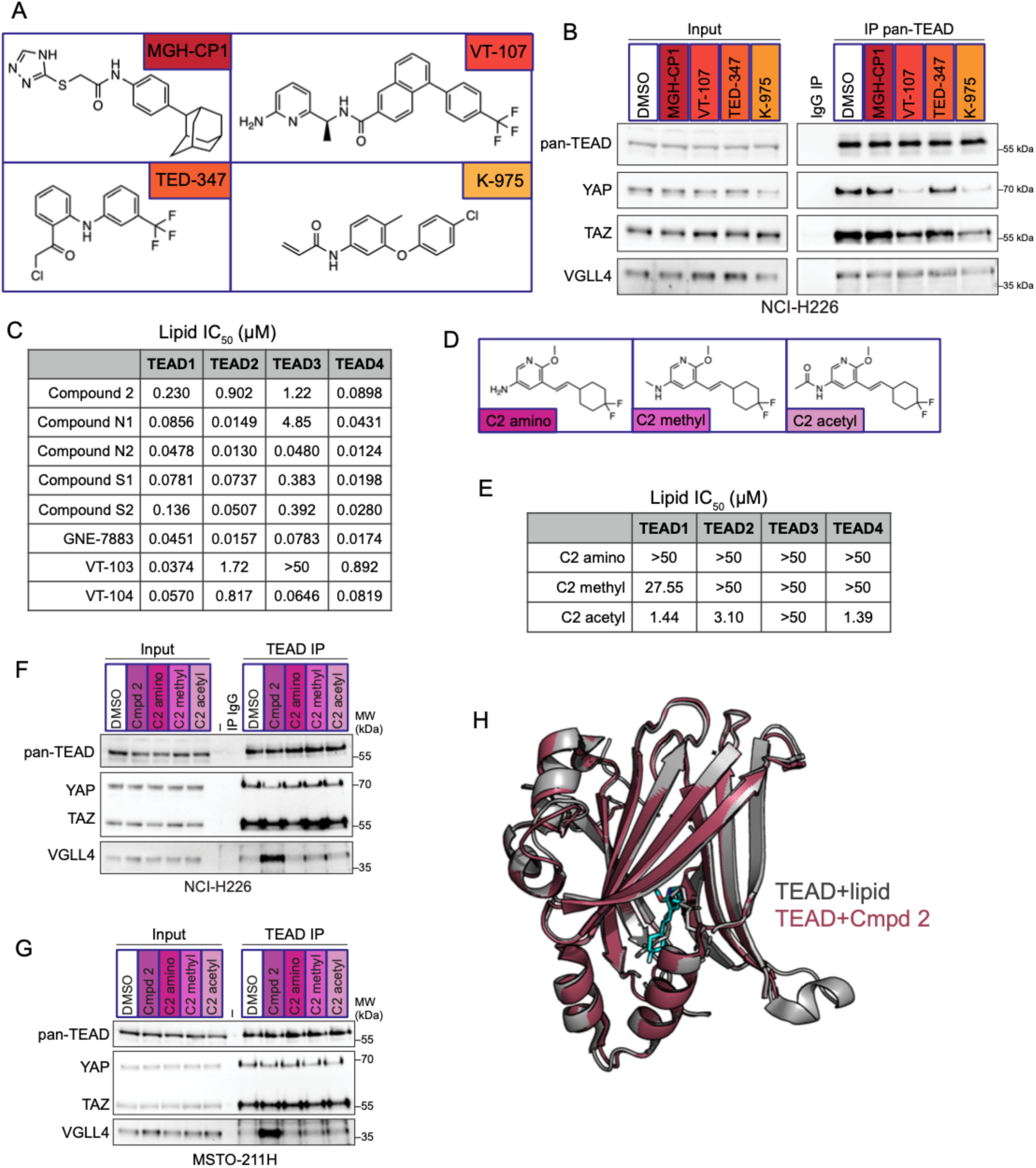
Profiling TEAD compounds and protein interactions. Related to Figure 2. (A) Structures of TEAD LPB molecules MGH-CP1, VT-107, TED-347, and K-975. (B) CoIP experiment to test TEAD compounds. NCI-H226cells were treated for 24 h with 3 µM of the indicated compounds or DMSO vehicle control and cellular extracts were subjected to IP with pan-TEAD antibody or an IgG control. CoIP samples were probed with antibodies against the indicated endogenous proteins. Inputs are 10% for pan-TEAD and 0.5% for YAP, TAZ, and VGLL4. n = 2 biological replicates. (C) Lipid displacement assay to measure compounds binding to the lipid pocket of TEAD using time-resolved fluorescence resonance energy transfer (TR-FRET). TEAD recombinant proteins are purified, 6xHis-tagged C-terminal domains for each TEAD: TEAD1(S210-D426), TEAD2(A217-D447), TEAD3(Q216-D435), and TEAD4(Q215-E434). (D) Structures of tool compounds designed with sulfonamide group substitutions. (E) Evaluation of compounds binding to the TEAD lipid pocket using TR-FRET, as in (C). (F) NCI-H226 cells were treated for 24 h with 3 µM of the indicated compounds or DMSO vehicle control and cellular extracts were subjected to IP with pan-TEAD antibody or an IgG control. IP samples were probed with antibodies against the indicated endogenous proteins. Inputs are 10% for pan-TEAD and 1% for YAP, TAZ, and VGLL4. n = 2 biological replicates. (G) MSTO-211H cells were treated for 24 h with 3 µM of the indicated compounds or DMSO vehicle control and cellular extracts were subjected to IP with pan-TEAD antibody or an IgG control. IP samples were probed with antibodies against the indicated endogenous proteins. Inputs are 10% for pan-TEAD and 1% for YAP, TAZ, and VGLL4. N = 2 biological replicates. (H) Overlay of cocrystal structures for TEAD2 with palmitoylation (PDB 5EMV) and with Compound 2 (PDB 6UYC). Compound 2 (cyan) overlaps with the palmitoyl lipid modification.

**Figure S3:**
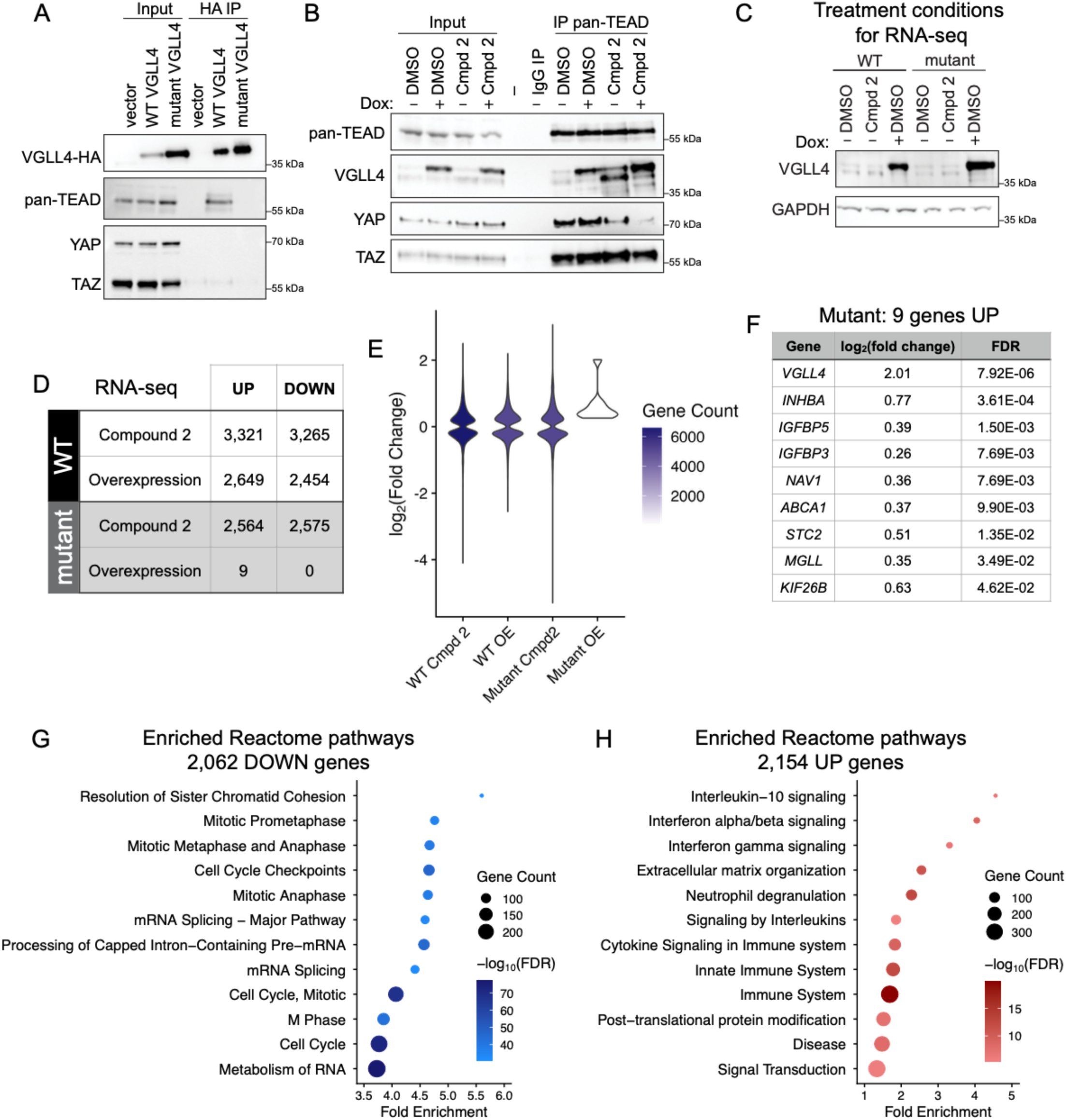
Inducible overexpression of VGLL4 enables transcriptional profiling. Related to Figure 3. (A) CoIP experiment to validate loss of TEAD interaction with mutant VGLL4 in NCI-H226 cells. VGLL4 expression was induced by 24 h doxycycline treatment and cellular extracts were subjected to IP with HA-conjugated beads. CoIP samples were probed with antibodies against the indicated endogenous proteins. Inputs are 20% for VGLL4 and 1% for TEAD, YAP, and TAZ. n = 3 biological replicates. (B) Pan-TEAD coIP experiment in NCI-H226 cells with Compound 2 treatment and with VGLL4 overexpression. Cells were treated for 24 h, IPs were performed on cellular extracts, and samples were probed with antibodies against the indicated endogenous proteins. Inputs are 10% for pan-TEAD and 1% for VGLL4, YAP, and TAZ. n = 1 biological replicate. (C) RNA-seq sample validation. Immunoblots of lysates from NCI-H226 cells expressing doxycycline-inducible VGLL4 constructs. (D) Number of transcripts significantly (FDR < 0.05) altered by 24 h treatment of cells with Compound 2 or doxycycline to induce overexpression of VGLL4, compared with DMSO control. n = 3 biological replicates. (E) Violin plot showing the distribution and magnitude (log_2_-fold) of the significant (FDR < 0.05) transcript changes elicited in the indicated conditions measured by RNA-seq. (F) Table listing the nine genes that change upon overexpression of mutant VGLL4. (G) Enrichment analysis on 2,062 commonly decreased genes from RNA-seq. Reactome Knowledgebase pathways were ranked by FDR and the top 12 pathways are presented. Color indicates the FDR, size indicates the number of genes. (H) Enrichment analysis on 2,154 commonly decreased genes from RNA-seq. Reactome Knowledgebase pathways were ranked by FDR and the top 12 pathways are presented. Color indicates the FDR, size indicates the number of genes.

**Figure S4:**
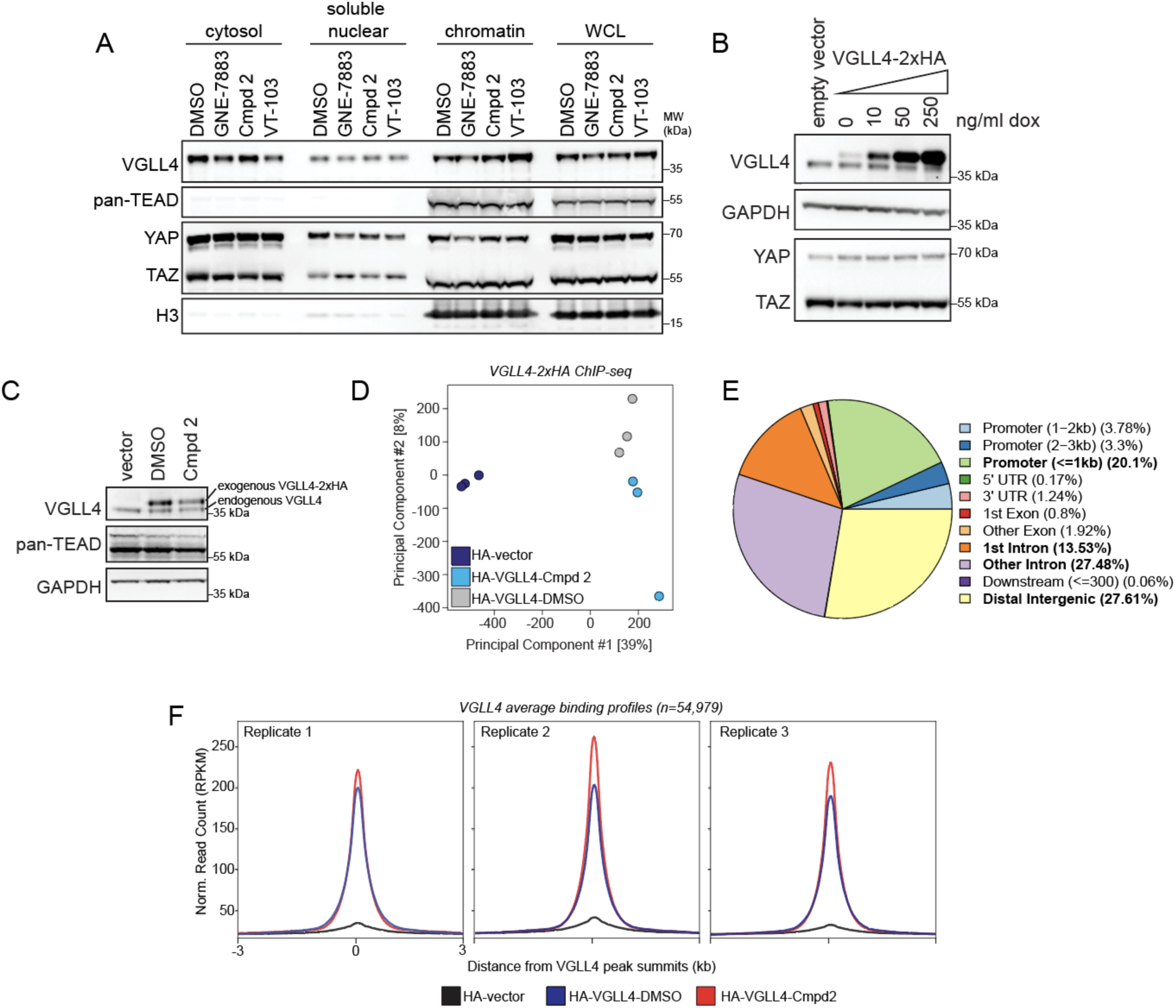
Inducible expression of VGLL4-2xHA enables genomic profiling. Related to Figure 4. (A) NCI-H226 cells were treated for 24 h and then fractionated into cytoplasmic, soluble nuclear, and chromatin-associated fractions. Equal amounts of each fraction were analyzed by immunoblotting with the antibodies against the indicated proteins. n = 4 biological replicates. (B) Titration of doxycycline (dox) in NCI-H226 cells for inducible exogenous expression of VGLL4-2xHA. (C) ChIP-seq sample validation. Immunoblots of lysates from NCI-H226 cells expressing doxycycline-inducible VGLL4 constructs and treated with 10 ng/ml doxycycline and compound. (D) PCA performed on ChIP-seq experimental samples. (E) Genomic feature annotation for VGLL4-HA ChIP-seq peaks. The four most representative categories are in bold. (F) Metagene plots of the HA-ChIP signal for each of the three biological replicates.

**Figure S5:**
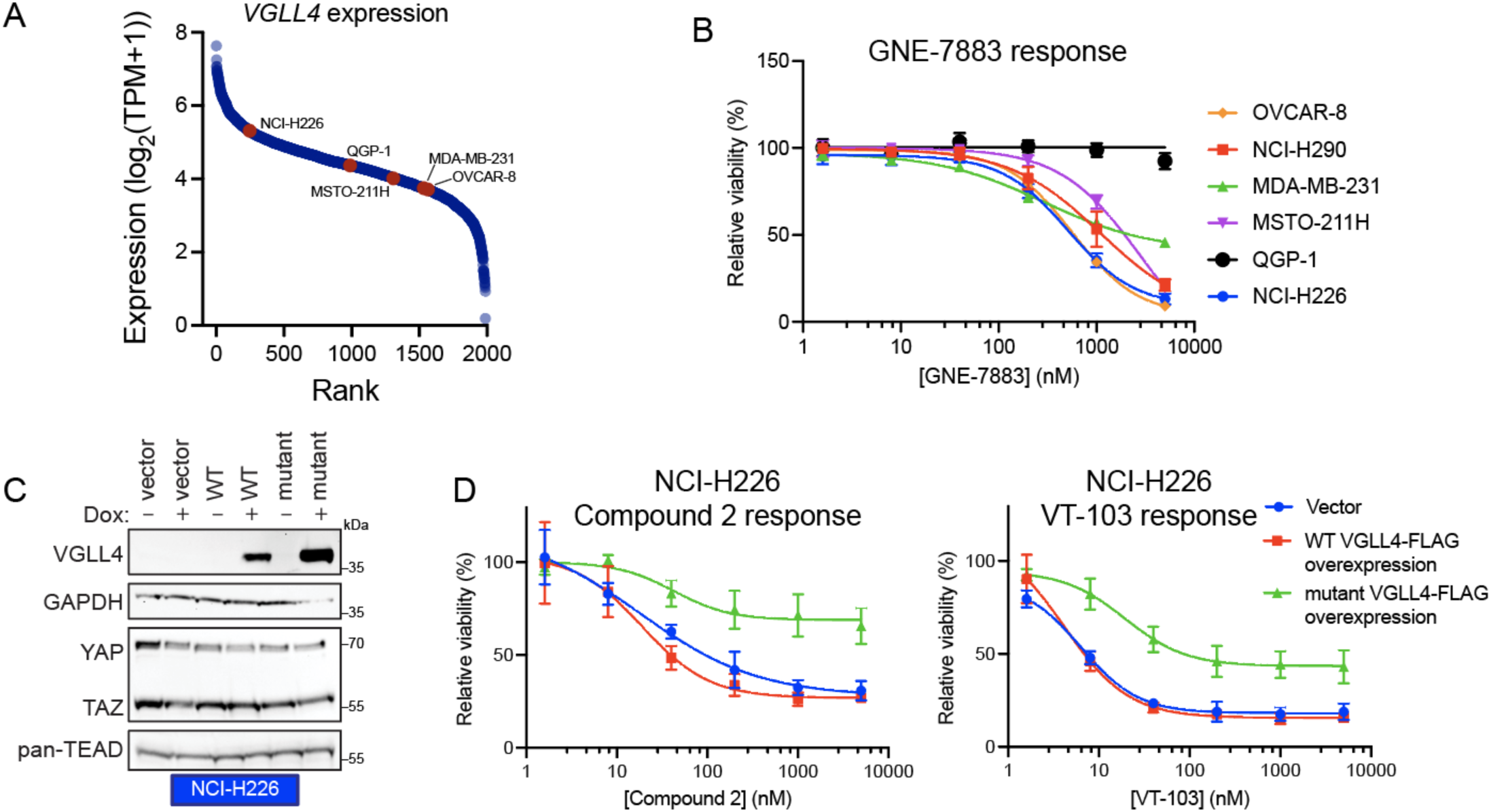
VGLL4 levels influence cell sensitivity to TEAD inhibitors. Related to Figure 5. (A) Rank of *VGLL4* RNA expression data from DepMap (Public 2024Q2) for the 1989 cell lines in DepMap. Specific cell lines investigated in this figure are highlighted in red. Note, the NCI-H290 cell line is not profiled in DepMap. TPM is transcripts per million. (B) Viability dose response curves (mean ± SD) for GNE-7883 in the six indicated cell lines treated for six days. n = 3 biological replicates. (C) Immunoblots of lysates from NCI-H226 cells engineered with inducible VGLL4 overexpression for the five indicated cell types treated with and without doxycycline (Dox) induction. The VGLL4 mutant used in these experiments is detailed in Figure 3A. (D) Viability dose response curves (mean ± SD) for Compound 2 (left) and VT-103 (right) in NCI-H226 cells. Viability assays were performed with uniform doxycycline treatment and five-fold titrated compounds. n = 3 biological replicates.

**Figure S6:**
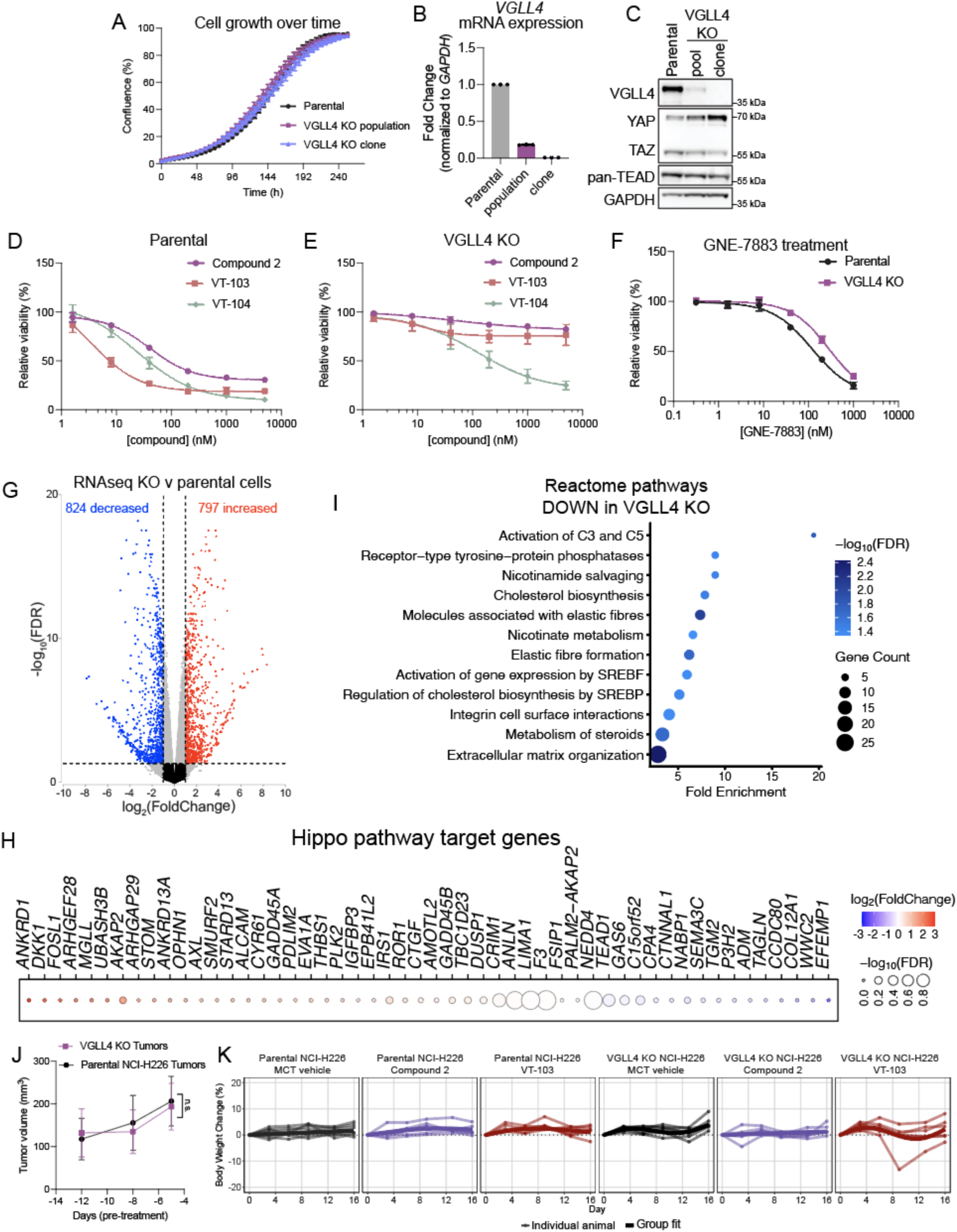
VGLL4 knockout confers resistance to Compound 2 and VT-103. Related to Figure 6. (A) Proliferation of NCI-H226 cells monitored over ten days. Data are represented as mean ± SEM; n = 3 biological replicates. (B) Analysis of *VGLL4* mRNA levels in NCI-H226 cells. Data are represented as mean ± SD; n = 3 biological replicates. (C) Immunoblots to analyze NCI-H226 VGLL4 knockout cells, pool and clone, after CRISPR genome editing. (D) Viability dose response curves (mean ± SD) for parental NCI-H226 cells treated for six days. n = 3 biological replicates. (E) Viability dose response curves (mean ± SD) for VGLL4 KO NCI-H226 cells treated for six days. n = 3 biological replicates. (F) Viability dose response curves (mean ± SD) for GNE-7883 treatment of parental and VGLL4 KO NCI-H226 cells treated for six days. n = 3 biological replicates. (G) Volcano plot depicting changes by RNA-seq in VGLL4 KO cells compared to parental NCI-H226 cells. Highly significant data points are colored red or blue and meet the cutoff FDR < 0.05 and |log_2_FoldChange| > 1. (H) Transcript level changes for 52 Hippo target genes upon VGLL4 knockout as measured by RNA-seq. FDR is represented by size and fold change is represented by color. (I) Pathway enrichment analysis on decreased genes from RNA-seq (FDR < 0.05 and |log_2_FoldChange| > 1). Reactome Knowledgebase pathways were ranked by FDR and the 12 enriched pathways are shown. Circle color indicates the FDR, size indicates the number of genes. (J) Tumor initiation growth measurements (mean ± SD) for parental NCI-H226 tumors and VGLL4 KO tumors measured before treatments. n.s. = not significant by unpaired t tests. n = 24 per genotype. (K) Percent changes in body weight of mice treated as indicated over time. The fitted averages are shown on top of the measurements for individual animals. n = 6 mice per group.

**Table S3:** Hippo gene signature of 52 genes. Related to Figures 2 and 6.

| **Gene Symbol** | **ENSEMBL ID** |
| --- | --- |
| *ADM* | ENSG00000148926 |
| *AKAP2* | ENSG00000241978 |
| *ALCAM* | ENSG00000170017 |
| *AMOTL2* | ENSG00000114019 |
| *ANKRD1* | ENSG00000148677 |
| *ANKRD13A* | ENSG00000076513 |
| *ANLN* | ENSG00000011426 |
| *ARHGAP29* | ENSG00000137962 |
| *ARHGEF28* | ENSG00000214944 |
| *AXL* | ENSG00000167601 |
| *C15orf52* | ENSG00000188549 |
| *CCDC80* | ENSG00000091986 |
| *COL12A1* | ENSG00000111799 |
| *CPA4* | ENSG00000128510 |
| *CRIM1* | ENSG00000150938 |
| *CTGF* | ENSG00000118523 |
| *CTNNAL1* | ENSG00000119326 |
| *CYR61* | ENSG00000142871 |
| *DKK1* | ENSG00000107984 |
| *DUSP1* | ENSG00000120129 |
| *EFEMP1* | ENSG00000115380 |
| *EPB41L2* | ENSG00000079819 |
| *EVA1A* | ENSG00000115363 |
| *F3* | ENSG00000117525 |
| *FOSL1* | ENSG00000175592 |
| *FSIP1* | ENSG00000150667 |
| *GADD45A* | ENSG00000116717 |
| *GADD45B* | ENSG00000099860 |
| *GAS6* | ENSG00000183087 |
| *IGFBP3* | ENSG00000146674 |
| *IRS1* | ENSG00000169047 |
| *LEPREL1* | ENSG00000090530 |
| *LIMA1* | ENSG00000050405 |
| *MGLL* | ENSG00000074416 |
| *NABP1* | ENSG00000173559 |
| *NEDD4* | ENSG00000069869 |
| *OPHN1* | ENSG00000079482 |
| *PALM2-AKAP2* | ENSG00000157654 |
| *PDLIM2* | ENSG00000120913 |
| *PLK2* | ENSG00000145632 |
| *ROR1* | ENSG00000185483 |
| *SEMA3C* | ENSG00000075223 |
| *SMURF2* | ENSG00000108854 |
| *STARD13* | ENSG00000133121 |
| *STOM* | ENSG00000148175 |
| *TAGLN* | ENSG00000149591 |
| *TBC1D23* | ENSG00000036054 |
| *TEAD1* | ENSG00000187079 |
| *TGM2* | ENSG00000198959 |
| *THBS1* | ENSG00000137801 |
| *UBASH3B* | ENSG00000154127 |
| *WWC2* | ENSG00000151718 |

