## Supplementary material for "TEAD-targeting small molecules induce a cofactor switch to regulate the Hippo pathway": Methods S1

**Contents**

General considerations ... 2
Synthesis of Compound S1 ... 3
Synthesis of Compound N1 ... 4
Synthesis of Compound S2 ... 9
Synthesis of Compound N2 ... 11
Synthesis of C2-Amino ... 14
Synthesis of C2-Acetyl ... 17
Synthesis of C2-Methyl ... 18

**General considerations**

All chemicals were used directly as received from commercial suppliers. $^{1}$H-NMR and $^{13}$C-NMR spectra were recorded on Bruker Avance 400 or 500 spectrometers in deuterated solvents (DMSO-$d_6$ or $CDCl_3$). Chemical shifts are expressed in δ ppm referenced to an internal standard, tetramethylsilane (δ = 0 ppm). Abbreviations used in describing peak signals are: br = broad signal, s = singlet, d = doublet, dd = doublet of doublets, t = triplet, q = quartet, m = multiplet. All final compounds were purified to > 95% by reverse phase high performance liquid chromatography (HPLC), super-critical fluid chromatography (SFC) or normal phase silica gel flash chromatography. The purity was assessed by reverse phase HPLC with a gradient of 5%–95% acetonitrile in water (with either acid or base modifier) and monitored by absorption at 254 nm. Low-resolution mass spectra were recorded on liquid chromatography-mass spectrometer in electrospray positive (ES+) mode. High-resolution mass spectrometry (HRMS) experiments were performed on Dionex LC Ultimate 3000 coupled with ThermoScientific Q Exactive orbitrap mass spectrometer using ESI as ionization source and a Phenomenex XB-C18, 1.7mm, 50 3 2.1 mm column with a 0.7 mL / minute flow rate at 40˚C for LC separation. Solvent A is water with 0.1% FA and solvent B is acetonitrile with 0.1% FA. The gradient consisted with 2 - 98% solvent B over 7 min and hold 98% B for 1.5 min following equilibration for 1.0 min. The LC was monitored by absorption at 220 nm and 254 nm. MS full scan with 10,000 resolutions was applied to all experiments.

**Synthesis of Compound S1**

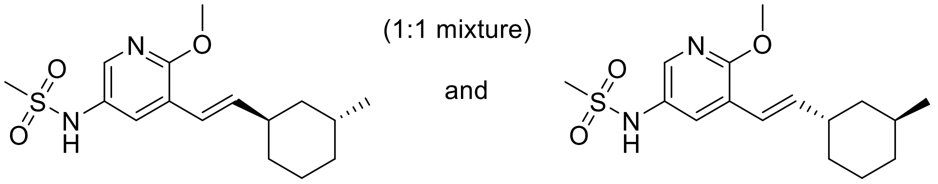

N-(6-methoxy-5-((E)-2-((1R,3R)-3-methylcyclohexyl)vinyl)pyridin-3-yl)methanesulfonamide and N-(6-methoxy-5-((E)-2-((1S,3S)-3-methylcyclohexyl)vinyl)pyridin-3-yl)methanesulfonamide (1:1 mixture)

**Step 1:** Preparation of (E)-5-bromo-2-methoxy-3-(2-(3-methylcyclohexyl)vinyl)pyridine

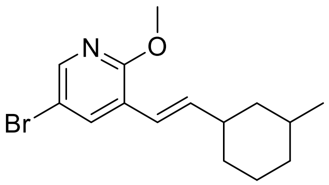

To a stirred mixture of 5-bromo-3-(diethoxyphosphorylmethyl)-2-methoxy-pyridine (1.5 g, 4.4 mmol) in anhydrous tetrahydrofuran (30 mL) cooled to 0 °C was added sodium hydride (60% mass) in mineral oil (200 mg, 4.9 mmol) and the reaction mixture was stirred at 0 °C for 1 h. A solution of 3-methylcyclohexanecarbaldehyde (620 mg, 4.9 mmol) in tetrahydrofuran was added. The reaction mixture was stirred at room temperature for 48 h, then quenched with saturated aqueous ammonium chloride solution and extracted with dichloromethane. The combined organic layers were dried over sodium sulfate, filtered, and concentrated under reduced pressure. The crude residue was purified by column chromatography (silica gel: isopropyl acetate / heptane) to give 387 mg (28% yield) of (E)-5-bromo-2-methoxy-3-(2-(3-methylcyclohexyl)vinyl)pyridine as a white solid. MS (ESI+) m/z 310 $(M+H)^+$.

**Step 2:** Preparation of S1 (1:1 mixture)

Into a vial was placed (E)-5-bromo-2-methoxy-3-(2-(3-methylcyclohexyl)vinyl)pyridine (390 mg, 1.30 mmol), methanesulfonamide (180 mg, 1.9 mmol), tris(dibenzylideneacetone)dipalladium(0) (88 mg, 0.09 mmol), 2-di-tert-butylphosphino-2',4',6'-triisopropylbiphenyl (164 mg, 0.37 mmol), potassium carbonate (430 mg, 3.1 mmol) and 2-methyltetrahyrofuran (2 mL). The reaction mixture was vacuum purged with nitrogen and heated to 100 °C for 14 h. The reaction was cooled to room temperature and then filtered through a pad of Celite eluted with dichloromethane. The filtrate was concentrated under reduced pressure. The crude residue was purified by reverse-phase HPLC, and the diasteromers were separated to collect 51 mg (13%) of the title compounds as 1:1 mixture of the trans-enantiomers.

**$^1$H NMR** (400 MHz, DMSO) δ 9.47 (s, 1H), 7.90 (d, J = 2.7 Hz, 1H), 7.62 (d, J = 2.6 Hz, 1H), 6.42 (d, J = 16.1 Hz, 1H), 6.28 (dd, J = 16.1, 6.7 Hz, 1H), 3.87 (s, 3H), 2.95 (s, 3H), 2.23-2.11 (m, 1H), 1.78-1.69 (m, 3H), 1.65 (d, J = 13.1 Hz, 1H), 1.50-1.36 (m, 1H), 1.35-1.27 (m, 1H), 1.10-0.97 (m, 1H), 0.89 (d, J = 6.5 Hz, 3H), 0.87-0.75 (m, 2H).

**$^{13}$C NMR** (101 MHz, DMSO) δ 158.03, 140.26, 139.17, 129.96, 129.29, 120.90, 120.51, 53.94, 41.44, 41.38, 34.77, 32.33, 25.93, 23.21.

**HRMS (ESI+)** m/z found $MH^+$ 325.1581, $C_{16}H_{25}N_2O_3S$ requires 325.1586.

**Synthesis of Compound N1**

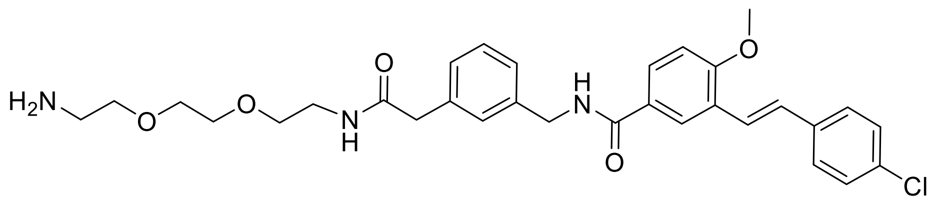

(E)-N-(3-(2-((2-(2-(2-aminoethoxy)ethoxy)ethyl)amino)-2-oxoethyl)benzyl)-3-(4-chlorostyryl)-4-methoxybenzamide

**Step 1:** Preparation of ethyl 3-formyl-4-hydroxybenzoate

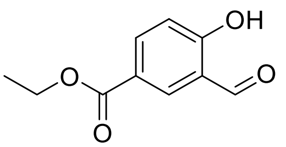

To a stirred solution of ethyl 4-hydroxybenzoate (100 g, 0.6 mol) and triethylamine (450 mL, 3.6 mol) in dichloromethane (1 L) was added magnesium chloride (285 g, 3.0 mol), and the reaction mixture was stirred at 40 °C for 1 h. Paraformaldehyde (180 g, 6.0 mol) was then added and the mixture was stirred at 70 °C for 3 h. After cooling to 0 °C, 1N hydrochloric acid (3 L) was slowly added. The solid was filtered and then washed with dichloromethane (0.17 L). The filtrate was washed with 1N hydrochloric acid (0.17 L) and brine (0.17 L). The organic phase was dried over magnesium sulfate, filtered, and concentrated under reduced pressure. The crude product was purified by column chromatography (silica gel: ethyl acetate / hexane) to afford 80 g (68% yield) of ethyl 3-formyl-4-hydroxybenzoate.

**$^1$H NMR** (300 MHz, $CDCl_3$) δ 11.40 (s, 1H), 9.97 (s, 1H), 8.34 (s, 1H), 8.21 (d, J = 10.8 Hz, 1H), 7.05 (d, J = 8.7 Hz, 1H), 4.41 (q, J = 7.2 Hz, 2H), 1.42 (t, J = 7.2 Hz, 3H).

**Step 2:** Preparation of ethyl 3-formyl-4-methoxybenzoate

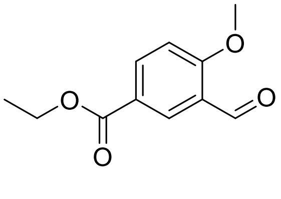


To a stirred solution of ethyl 3-formyl-4-hydroxybenzoate (155 g, 0.8 mol) in acetone (1.5 L) was added potassium carbonate (144 g, 1.1 mol) and dimethyl carbonate (86.4 g, 1.0 mol). The reaction mixture was stirred at reflux for 1 h and then cooled to room temperature. The resultant solid was filtered and washed with ethyl acetate (3 x 0.1 L). The filtrate was diluted with ethyl acetate (1 L) and saturated aqueous sodium bicarbonate solution (1 L). The biphasic layers were separated, and the aqueous layer was further extracted with ethyl acetate (3 x 1 L). The combined organic layers were washed with water (2 x 1 L), dried over magnesium sulfate, and concentrated under reduced pressure. The crude product was purified by column chromatography (silica gel: ethyl acetate / hexane) to afford 115 g (69% yield) of ethyl 3-formyl-4-methoxybenzoate.

**$^1$H NMR** (300 MHz, $CDCl_3$) δ 10.47 (s, 1H), 8.51 (s, 1H), 8.26 (d, J = 10.8 Hz, 1H), 7.05 (d, J = 8.7 Hz, 1H), 4.38 (q, J = 7.2 Hz, 2H), 4.02 (s, 3H), 1.41 (t, J = 7.2 Hz, 3H).

**Step 3:** Preparation of ethyl (E)-3-(4-chlorostyryl)-4-methoxybenzoate

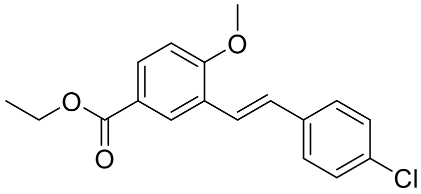


To a solution of diethyl 4-chlorobenzylphosphonate (160 g, 0.77 mol) in toluene (1.5 L) at 0 °C was added sodium tert-pentoxide (89 g, 0.87 mol), and the reaction mixture was stirred at 0 °C for 20 min. To this mixture was added dropwise a solution of ethyl 3-formyl-4-methoxybenzoate

(120 g, 0.58 mol) in tetrahydrofuran (0.5 L) over 20 min. The reaction mixture was stirred for 1.5 h at 0 °C and quenched with saturated aqueous ammonium chloride solution (3 L). The mixture was extracted with ethyl acetate (2 x 2 L). The combined organic layers were washed with brine (2 L), dried over sodium sulfate, filtered, and concentrated under reduced pressure to give 160 g (87% yield) of crude ethyl (E)-3-(4-chlorostyryl)-4-methoxybenzoate, which was used in the next reaction step without further purification.

**Step 4:** Preparation of (E)-3-(4-chlorostyryl)-4-methoxybenzoic acid

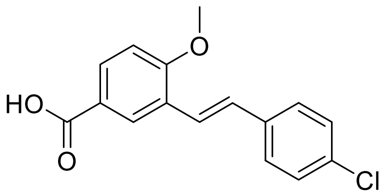


To a solution of ethyl (E)-3-(4-chlorostyryl)-4-methoxybenzoate (160 g, 0.50 mol) in methanol (1 L) was added aqueous potassium hydroxide (20 mass %) solution (260 mL). The mixture was stirred at 65 °C for 2 h and then cooled to 0 °C. The reaction mixture was adjusted to pH ~ 3 by the addition of 1N hydrochloric acid. The resulting precipitate was filtered to afford 104 g (72% yield) of (E)-3-(4-chlorostyryl)-4-methoxybenzoic acid as a white solid.

**$^1$H NMR** (300 MHz, DMSO) δ 12.77 (br s, 1H), 8.21 (s, 1H), 7.89 (d, J = 8.7 Hz, 1H), 7.66-7.63 (m, 2H), 7.46-7.41 (m, 3H), 7.27-7.17 (m, 1H), 7.15 (d, J = 8.7 Hz, 1H), 3.94 (s, 3H).

**MS (ESI-)** m/z 287 $(M-H)^-$.

**Step 5:** Preparation of methyl (E)-2-(4-((3-(4-chlorostyryl)-4-methoxybenzamido)methyl)-phenyl)acetate

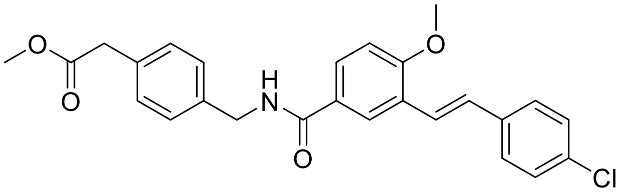


A mixture of (E)-3-(4-chlorostyryl)-4-methoxybenzoic acid (630 mg, 2.1 mmol), methyl 2-[3-(aminomethyl)phenyl]acetate hydrochloride (610 mg, 2.8 mmol), hexafluorophosphate azabenzotriazole tetramethyl uronium (1200 mg, 3.3 mmol), and diisopropylethylamine (1.5 mL, 8.7 mmol) in N,N-dimethylformamide (7.3 mL) was stirred at room temperature. After 3 days the reaction was diluted with isopropyl acetate. The organic layer was washed with water and brine, dried over sodium sulfate, filtered, and concentrated under reduced pressure. The crude product was purified by column chromatography (silica gel: methanol / isopropyl acetate) to afford 880 mg (90% yield) of methyl (E)-2-(4-((3-(4-chlorostyryl)-4-methoxybenzamido)methyl)-phenyl)acetate as a white foam.

**$^1$H NMR** (400 MHz, $CDCl_3$) δ 8.05 (d, J = 2.3 Hz, 1H), 7.69 (dd, J = 8.5, 2.3 Hz, 1H), 7.48-7.43 (m, 2H), 7.40 (d, J = 16.6 Hz, 1H), 7.36-7.27 (m, 5H), 7.24-7.20 (m, 1H), 7.14 (d, J = 16.5 Hz, 1H), 6.91 (d, J = 8.6 Hz, 1H), 6.41 (t, J = 5.8 Hz, 1H), 4.65 (d, J = 5.7 Hz, 2H), 3.93 (s, 3H), 3.69 (s, 3H), 3.63 (s, 2H).

**MS (ESI+)** m/z 450 $(M+H)^+$.

**Step 6:** Preparation of (E)-2-(4-((3-(4-chlorostyryl)-4-methoxybenzamido)methyl)phenyl)acetic acid

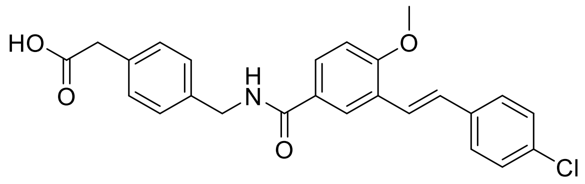


To a mixture of methyl (E)-2-(4-((3-(4-chlorostyryl)-4-methoxybenzamido)methyl)-phenyl)acetate (860 mg, 1.9 mmol) dissolved in tetrahydrofuran (9.6 mL), methanol (3.2 mL), and water (3.2 mL) was added lithium hydroxide (0.08 mL), and the reaction mixture was stirred at 40 °C for 14h. Volatile solvent was reduced under pressure, and the reaction mixture was acidified with 1N hydrochloric acid until pH ~ 1 and then diluted with isopropyl acetate. The layers were separated and the aqueous layer was further washed with isopropyl acetate (3x). The combined organic layers were washed with water, dried over sodium sulfate, filtered, and concentrated under reduced pressure. The crude product was triturated with dichloromethane / hexane to give 760 mg (91% yield) of (E)-2-(4-((3-(4-chlorostyryl)-4-methoxybenzamido)methyl)phenyl)acetic acid as a white solid.

**$^1$H NMR** (400 MHz, DMSO) δ 8.98 (t, J = 6.0 Hz, 1H), 8.24 (d, J = 2.3 Hz, 1H), 7.87 (dd, J = 8.6, 2.2 Hz, 1H), 7.61 (d, J = 8.6 Hz, 2H), 7.47-7.38 (m, 3H), 7.29 (d, J = 16.5 Hz, 1H), 7.24-7.16 (m, 2H), 7.15-7.07 (m, 3H), 4.46 (d, J = 5.9 Hz, 2H), 3.91 (s, 3H), 3.37 (s, 2H); $CO_2H$ not seen.

**MS (ESI+)** m/z 436 $(M+H)^+$.

**Step 7:** Preparation of tert-butyl (E)-(2-(2-(2-(2-(4-((3-(4-chlorostyryl)-4-methoxybenzamido)methyl)phenyl)acetamido)ethoxy)ethoxy)ethyl)carbamate

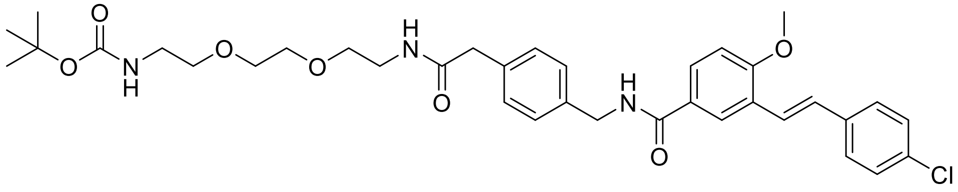


A mixture of (E)-2-(4-((3-(4-chlorostyryl)-4-methoxybenzamido)methyl)phenyl)acetic acid (460 mg, 1.1 mmol), tert-butyl N-[2-[2-(2-aminoethoxy)ethoxy]ethyl]carbamate (390 mg, 1.6 mmol), hexafluorophosphate azabenzotriazole tetramethyl uronium (680 mg, 1.8 mmol), and diisopropylethylamine (0.73 mL, 4.2 mmol) in N,N-dimethylformamide (3.5 mL) was stirred at room temperature for 42 h. The reaction mixture was diluted with isopropyl acetate and then washed with water and brine. The organic layer was dried over sodium sulfate, filtered, and concentrated under reduced pressure. The crude product was purified by column chromatography (silica: methanol / isopropyl acetate) to afford 678 mg (97% yield) of tert-butyl (E)-(2-(2-(2-(2-(4-((3-(4-chlorostyryl)-4-methoxybenzamido)methyl)phenyl)acetamido)ethoxy)ethoxy)ethyl)-carbamate.

**$^1$H NMR** (400 MHz, $CDCl_3$) δ 8.13 (d, J = 2.4 Hz, 1H), 7.78 (dd, J = 8.8, 2.4 Hz, 1H), 7.48-7.43 (m, 2H), 7.40 (d, J = 16.5 Hz, 1H), 7.33-7.23 (m, 7H), 7.15 (d, J = 16.5 Hz, 2H), 6.91 (d, J = 8.7 Hz, 1H), 6.31 (s, 1H), 5.14 (d, J = 6.9 Hz, 1H), 4.61 (d, J = 5.8 Hz, 2H), 3.92 (s, 3H), 3.53 (s, 2H), 3.50-3.43 (m, 7H), 3.39 (q, J = 5.2 Hz, 2H), 3.27-3.19 (m, 2H), 1.41 (s, 9H).

**MS (ESI+)** m/z 666 $(M+H)^+$.

**Step 8:** Preparation of S1

To a mixture of methyl tert-butyl (E)-(2-(2-(2-(2-(4-((3-(4-chlorostyryl)-4-methoxybenzamido)methyl)phenyl)acetamido)ethoxy)ethoxy)ethyl)-carbamate (680 mg, 1.0 mmol) in anhydrous methanol (3.4 mL) and dichloromethane (3.4 mL) was added a solution of hydrochloric acid (4 mol/L) in 1,4-dioxane (2.5 mL), and the reaction mixture was stirred at room temperature for 20h. Volatile solvent was removed under reduced pressure, and the reaction mixture was basified with aqueous sodium hydroxide solution (1N) until neutral pH. The mixture was washed with isopropyl acetate (5 x 10 mL). The combined organic layers were dried over sodium sulfate, filtered, and concentrated under reduced pressure. Trituration with dichloromethane resulted in a white solid which was filtered. The filtrate was concentrated under reduced pressure and purified by reverse-phase HPLC to give 49 mg (62% yield) of the title compound as a white solid.

**$^{1}$H NMR** (400 MHz, $CDCl_3$) δ 8.11 (d, J = 2.2 Hz, 1H), 7.75 (dd, J = 8.7, 2.3 Hz, 1H), 7.46-7.35 (m, 4H), 7.32-7.22 (m, 5H), 7.19-7.08 (m, 2H), 6.88 (d, J = 8.7 Hz, 1H), 6.60 (t, J = 5.6 Hz, 1H), 4.58 (d, J = 5.8 Hz, 2H), 3.90 (s, 3H), 3.76-3.73 (m, 1H), 3.67-3.60 (m, 1H), 3.51 (s, 2H), 3.50-3.41 (m, 5H), 3.37 (q, J = 5.3 Hz, 2H), 2.85-2.80 (m, 2H), 2.81-2.72 (m, 3H).

**$^{13}$C NMR** (101 MHz, DMSO) δ 170.16, 165.57, 158.68, 139.74, 136.45, 136.19, 131.99, 128.77, 128.66, 128.28, 128.10, 127.83, 127.42, 126.59, 125.53, 125.29, 124.83,123.34, 111.05, 70.54, 69.54, 69.50, 69.04, 55.90, 42.54, 42.23, 40.24, 38.67.

**HRMS (ESI+)** m/z found $MH^+$ 566.2414, $C_{31}H_{37}ClN_3O_5$ requires 566.2416.

**Synthesis of Compound S2**

(E)-N-(5-(2-(3,3-dimethylcyclohexyl)vinyl)-6-methoxypyridin-3-yl)methanesulfonamide – single unknown isomer*

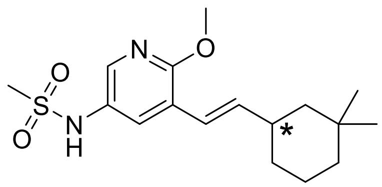

**Step 1:** Preparation of (E)-5-bromo-3-(2-(3,3-dimethylcyclohexyl)vinyl)-2-methoxypyridine

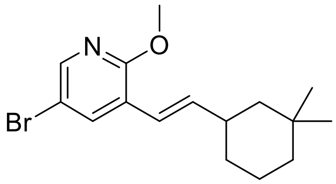

To a stirred solution of 5-bromo-3-(diethoxyphosphorylmethyl)-2-methoxy-pyridine (1.5 g, 4.4 mmol) in tetrahydrofuran (30 mL) at 0 °C was added sodium hydride (60% mass) in mineral oil (200 mg, 4.9 mmol), and the reaction mixture was stirred at 0 °C for 1 h. A solution of 3,3-dimethylcyclohexanecarbaldehyde (680 mg, 4.9 mmol) in tetrahydrofuran was added. The reaction mixture was stirred at room temperature for 48 h, then quenched with saturated aqueous ammonium chloride solution and extracted with dichloromethane. The combined organic layers were dried over sodium sulfate, filtered, and concentrated under reduced pressure. The crude was purified by column chromatography (silica gel: isopropyl acetate / heptane) to give 540 mg (38% yield) of (E)-5-bromo-3-(2-(3,3-dimethylcyclohexyl)vinyl)-2-methoxypyridine as a colorless oil.

**MS (ESI+)** m/z 324 $(M+H)^+$.

**Step 2:** Preparation of S2

To a vial was placed (E)-5-bromo-3-(2-(3,3-dimethylcyclohexyl)vinyl)-2-methoxypyridine (540 mg, 1.67 mmol), methanesulfonamide (240 mg, 2.5 mmol), tris(dibenzylideneacetone)dipalladium(0) (118 mg, 0.13 mmol), 2-di-tert-butylphosphino-2',4',6'-triisopropylbiphenyl (220 mg, 0.50 mmol), potassium carbonate (580 mg, 4.2 mmol) and 2-methyltetrahyrofuran (2.8 mL). The reaction mixture was vacuum purged with nitrogen and heated to 100 °C for 14 h. The reaction was cooled to room temperature and then filtered through a pad of Celite eluted with dichloromethane. The filtrate was concentrated under reduced pressure. The crude residue was purified by chiral SFC (Amylose-1: 0.1% Ammonium hydroxide in methanol). The first eluted peak was collected to give 44 mg (7.8% yield) of (E)-N-(5-(2-(3,3-dimethylcyclohexyl)vinyl)-6-methoxypyridin-3-yl)methanesulfonamide – single unknown isomer*.

**$^1$H NMR** (400 MHz, DMSO) δ 9.47 (s, 1H), 7.90 (d, J = 2.5 Hz, 1H), 7.62 (d, J = 2.5 Hz, 1H), 6.43 (d, J = 16.1 Hz, 1H), 6.25 (dd, J = 16.1, 6.8 Hz, 1H), 3.88 (s, 3H), 2.96 (s, 3H), 2.39-2.27 (m, 1H), 1.79-1.70 (m, 1H), 1.62-1.39 (m, 3H), 1.39-1.31 (m, 1H), 1.15-0.95 (m, 3H), 0.93 (d, J = 4.5 Hz, 6H).

**$^{13}$C NMR** (101 MHz, DMSO) δ 157.53, 140.03, 138.69, 129.48, 128.81, 120.41, 120.12, 53.45, 45.19, 38.36, 36.82, 33.19, 32.13, 30.41, 24.48, 21.53.

**HRMS (ESI+)** m/z found $MH^+$ 339.1739, $C_{17}H_{27}N_2O_3S$ requires 339.1742.

**Synthesis of Compound N2**

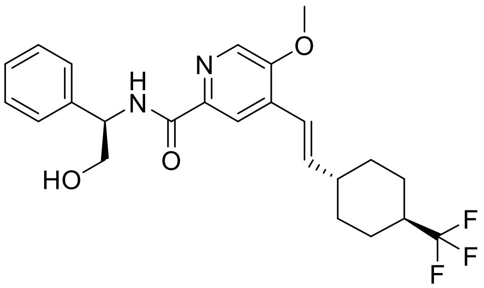

N-((R)-2-hydroxy-1-phenylethyl)-5-methoxy-4-((E)-2-(4-(trifluoromethyl)cyclohexyl)-vinyl)picolinamide

**Step 1:** Preparation of 4-(trifluoromethyl)cyclohexane-1-carbaldehyde

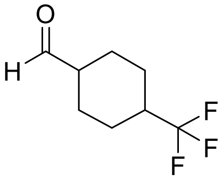

To a stirred solution of methyl 4-(trifluoromethyl)cyclohexanecarboxylate (10 g, 47.6 mmol in dichloromethane (0.15 M) cooled to -78 °C was added diisobutylaluminum hydride (1.0 mol/L) in tetrahydrofuran (50 mL). The reaction was then monitored by proton NMR. Upon completion the reaction mixture was quenched with methanol (30ml) followed by saturated aqueous Rochelle salt solution (500 mL). The reaction mixture was diluted with isopropyl acetate (2L). The biphasic layers were separated and the aqueous phase was further extracted with isopropyl acetate (4 x 1L). The combined organic layers were dried over magnesium sulfate, filtered, and concentrated under reduced pressure to give 8.6 g (100% yield) of 4-(trifluoromethyl)cyclohexane-1-carbaldehyde as a colorless oil, which was used immediately in the next reaction step.

**Step 2:** Preparation of (E)-2-chloro-5-methoxy-4-(2-(4-(trifluoromethyl)cyclohexyl)-vinyl) pyridine

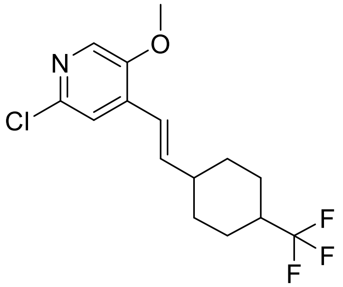


To a solution of 2-chloro-4-(diethoxyphosphorylmethyl)-5-methoxy-pyridine (1.0 g, 3.15 mmol) in toluene (18 mL) at 0 °C was added sodium tert-pentoxide (0.42 g, 3.78 mmol) and the mixture was stirred for 20 min at 0 °C. A solution of 4-(trifluoromethyl)cyclohexane-carbaldehyde (1.62 g, 6.3 mmol, 70% purity) in tetrahydrofuran (18 mL) was then added dropwise, and the reaction mixture was stirred at 0 °C for 1.5 h. The reaction mixture was poured into saturated aqueous ammonium chloride solution (50 mL) and extracted with ethyl acetate (2 x 100 mL). The organic layers were combined, washed with brine (50 mL), dried over sodium sulfate and concentrated under reduced pressure. The residue was purified by column chromatography on silica gel (ethyl acetate in petroleum ether) to afford 0.34 g (34% yield) of (E)-2-Chloro-5-methoxy-4-(2-(4-(trifluoromethyl)cyclohexyl)vinyl) pyridine as a colorless oil. MS (ESI+) m/z 320 $(M+H)^+$.

**Step 3:** Preparation of (E)-methyl 5-methoxy-4-(2-(4-(trifluoromethyl)cyclohexyl)vinyl) picolinate

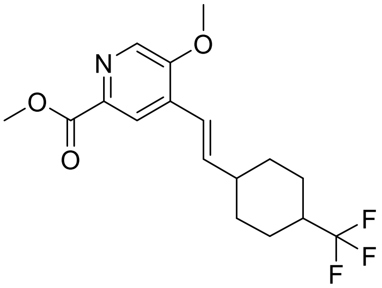


A mixture of (E)-2-Chloro-5-methoxy-4-(2-(4-(trifluoromethyl)cyclohexyl)vinyl) pyridine (0.32 g, 1 mmol), potassium carbonate (0.28 g, 2 mmol), palladium acetate (23 mg, 0.10 mmol) and l, 3-bis(diphenylphosphino)propane (83 mg, 0.20 mmol) in methanol (10 mL) and N,N-dimethylformamide (10 mL) was heated at 80 °C. under carbon monoxide atmosphere (50 psi) for 16 h. The reaction mixture was extracted with ethyl acetate (2 x30 mL) and washed with water (30 mL). The organic layer was dried over sodium sulfate and concentrated under reduced pressure. The crude residue was purified by prep-thin layer chromatography eluted with ethyl acetate in petroleum ether to afford 0.31 g (90% yield) of (E)-methyl 5-methoxy-4-(2-(4-(trifluoromethyl)cyclohexyl)-vinyl) picolinate as a light yellow oil. MS (ESI+) m/z 344 $(M+H)^+$.

**Step 4:** Preparation of (E)-5-methoxy-4-(2-(4-(trifluoromethyl)cyclohexyl)vinyl)picolinic acid

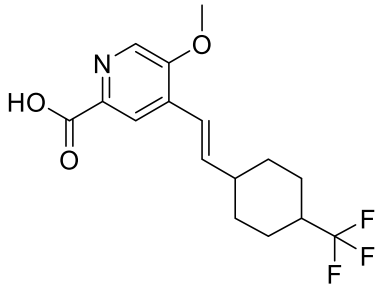

A mixture of lithium hydroxide hydrate (190 mg, 4.5 mmol) and (E)-methyl 5-methoxy-4-(2-(4-(trifluoromethyl)cyclohexyl)vinyl) picolinate (0.31 g, 0.90 mmol) in water (15 mL), methanol (15 mL), and tetrahydrofuran (3 mL) was stirred at 15 °C for 16 hours. The reaction mixture was concentrated under reduced pressure to remove organic solvent. The aqueous phase was adjusted to pH ~ 6 with aqueous 1 N hydrochloric acid and then extracted with ethyl acetate (2 x 30 mL). The combined organic layers were washed with water (30 mL), dried over sodium sulfate and concentrated under reduced pressure to give 200 mg (67% yield) of (E)-5-methoxy-4-(2-(4-(trifluoromethyl)cyclohexyl)vinyl)picolinic acid as a white solid. MS (ESI+) m/z 330 $(M+H)^+$.

**Step 5:** Preparation of N2

A mixture of 5-methoxy-4-((E)-2-(4-(trifluoromethyl)cyclohexyl)vinyl)picolinic acid (150 mg, 0.46 mmol), (2R)-2-amino-2-phenyl-ethanol (94 mg, 0.68 mmol), benzotriazol-1-yloxytripyrrolidinophosphonium hexafluorophosphate (296 mg, 0.57 mmol), and diisopropylethylamine (0.4 mL, 2.3 mmol) in N,N-dimethylformamide (1.8 mL) was stirred at room temperature for 8 h. The reaction mixture was diluted with isopropyl acetate and then washed with water and brine. The organic layer was dried over magnesium sulfate, filtered, and concentrated under reduced pressure. The crude residue was purified by chiral SFC (Chiralpak IC: 0.1% Ammonium hydroxide in methanol) to give 19.4 mg (9.5% yield) of the title compound as a white solid.

**$^1$H NMR** (400 MHz, DMSO) δ 8.82 (d, J = 8.3 Hz, 1H), 8.39 (s, 1H), 8.02 (s, 1H), 7.37 (d, J = 7.1 Hz, 2H), 7.31 (t, J = 7.4 Hz, 2H), 7.23 (t, J = 7.2 Hz, 1H), 6.66-6.51 (m, 2H), 5.08-5.99 (m, 2H), 4.01 (s, 3H), 3.81-3.71 (m, 2H), 2.33-2.15 (m, 2H), 1.98-1.82 (m, 4H), 1.43-1.19 (m, 4H).

**$^{13}$C NMR** (101 MHz, DMSO) δ 163.80, 154.13, 143.45, 142.17, 141.67, 133.86, 133.37, 132.74, 129.97, 128.58, 127.32, 127.29, 120.14, 118.60, 64.83, 57.12, 55.44, 40.32, 30.39, 24.57.

**HRMS (ESI+)** m/z found $MH^+$ 449.2045, $C_{24}H_{28}F_3N_2O_3$ requires 449.2052.

**Synthesis of C2-Amino**

(E)-5-(2-(4,4-difluorocyclohexyl)vinyl)-6-methoxypyridin-3-amine

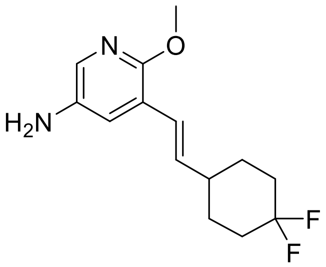

**Step 1:** Preparation of 5-bromo-2-methoxy-3-methylpyridine

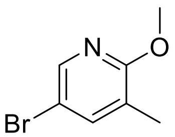

To a solution of 5-bromo-2-chloro-3-methylpyridine (300 g, 1.45 mol) in metanol (3 L) was added freshly prepared sodium methoxide (156 g, 2.9 mol), and the reaction mixture was heated to reflux and stirred overnight. The reaction mixture was quenched with acetic acid (600 mL) and concentrated under reduced pressure. The crude mixture was diluted with ethyl acetate (3 L) and washed with water (3 L). The aqueous layer was extracted with ethyl acetate (3 L), The combined organic layers were washed with brine (3 L), dried over anhydrous magnesium sulfate, and evaporated under reduced pressure to provide crude 5-bromo-2-methoxy-3-methylpyridine (220 g, 75% yield).

**$^1$H NMR** (300 MHz, $CDCl_3$) δ (s, 1 H), 7.47 (s, 1 H), 3.93 (s, 3 H), 2.05 (s, 3 H).

**Step 2:** Preparation of 5-bromo-3-(bromomethyl)-2-methoxypyridine

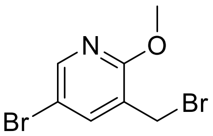

To a solution of 5-bromo-2-methoxy-3-methylpyridine (40 g, 198 mmol) in carbon tetrachloride (400 mL) was added N-bromosuccinimide (38.7 g, 217.4 mmol) and azobisisobutyronitrile (1.62 g, 6.1 mmol), and the reaction mixture was heated to reflux and stirred for 2 h. The reaction

mixture was concentrated under reduced pressure to give a crude residue. Petroleum ether (800 mL) was added, and the resultant solid was filtered. The filtrate was concentrated under reduced pressure to give a crude residue, which was triturated in petroleum ether to afford 5-bromo-3-(bromomethyl)-2-methoxypyridine as off-white solid (22 g, 40% yield). $^1$H NMR (300 MHz, $CDCl_3$) δ (s, 1 H), 7.73 (s, 1 H), 4.43 (s, 2 H), 4.00 (s, 3 H).

**Step 3:** Preparation of 5-bromo-3-(diethoxyphosphorylmethyl)-2-methoxypyridine

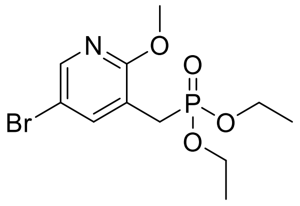

To a solution of 5-bromo-3-(bromomethyl)-2-methoxypyridine (22 g, 78.5 mmol) in 1,4-dioxane (110 mL) was added triethyl phosphite (26 g, 217.4 mmol), and the reaction mixture was heated to reflux and stirred overnight. The reaction mixture was concentrated under reduced pressure to remove volatile solvent, and the product was distilled to afford 5-bromo-3-(diethoxyphosphorylmethyl)-2-methoxypyridine as colorless oil (25 g, 94% yield).

**$^1$H NMR** (300 MHz, DMSO-$d_6$) δ (s, 1 H), 7.70 (s, 1 H), 4.09 (q, J = 7.2 Hz, 4 H), 3.94 (s, 3 H), 3.15 (d, J = 21.9 Hz, 2 H), 1.27 (t, J = 7.2 Hz, 6 H). MS (ESI+) m/z 337.8 $(M+H)^+$.

**Step 4:** Preparation of (E)-5-bromo-3-(2-(4.4-difluorocyclohexyl)vinyl)-2-methoxypyridine

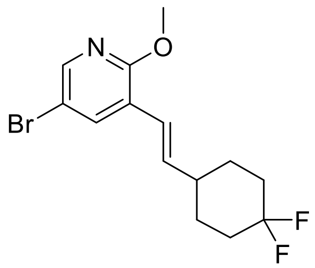

To a mixture of 4,4-difluorocyclohexanecarbaldehyde (1070 mg, 7.23 mmol) and 5-bromo-3-(diethoxyphosphorylmethyl)-2-methoxypyridine (820 mg, 2.41 mmol) in tetrahydrofuran (13.4 mL) was added potassium tert-butoxide (1910 mg, 16.9 mmol), and the reaction mixture was stirred at room temperature under nitrogen atmosphere for 2 h. The reaction mixture was diluted with isopropyl acetate and water. The organic phase was washed with water and brine, dried over sodium sulfate, filtered, and concentrated under reduced pressure. The crude residue was purified by column chromatography (silica gel: isopropyl acetate / heptane) to obtain (E)-5-bromo-3-(2-(4,4-difluorocyclohexyl)vinyl)-2-methoxypyridine (258 mg, 32% yield).

**MS (ESI+)** m/z 332/334 $(M+H)^+$.

**Step 5:** Preparation of (E)-N-(5-(2-(4,4-difluorocyclohexyl)vinyl)-6-methoxypyridin-3-yl)-l,l-diphenylmethanimine

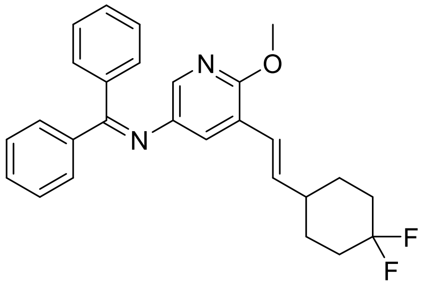


In a 20-mL vial was placed 5-bromo-3-[(E)-2-(4,4-difluorocyclohexyl)vinyl]-2-methoxy-pyridine (257 mg, 0.8 mmol), diphenylmethanimine (0.18 mL, 1.08 mmol), sodium tert-butoxide (149 mg, 1.6 mmol), bis(2-diphenylphosphinophenyl)ether (42 mg, 0.08 mmol), and tris(dibenzylidenteactone)dipalladium(0) (35 mg, 0.04 mmol). Degassed toluene (5.2 mL) was added. The vial was vacuum purged and back-filled with nitrogen (3x) and capped. The reaction mixture was stirred at 120 °C for 40 h. The reaction mixture was diluted with isopropyl acetate and water and then filtered through a pad of Celite. The biphasic layers were separated. The organic phase was washed with water and brine, dried over sodium sulfate, filtered, and concentrated under reduced pressure. The crude residue was purified by column chromatography (silica: isopropyl acetate / heptane) to obtain (E)-N-(5-(2-(4,4-difluorocyclohexyl)vinyl)-6-methoxypyridin-3-yl)-l,l-diphenylmethanimine (165 mg, 49% yield) as a yellow oil.

**MS (ESI+)** m/z 433 $(M+H)^+$.

**Step 6:** Preparation of C2-Amino
To (E)-N-(5-(2-(4,4-difluorocyclohexyl)vinyl)-6-methoxypyridin-3-yl)-l,l-diphenylmethanimine (165 mg, 0.38 mmol) dissolved in tetrahydrofuran (7.7 mL) was added aqueous hydrochloric acid (3.8 mL, 3.87 mmol, 1N), and the reaction mixture was stirred at room temperature for 2 h. Volatile solvent was removed under reduced pressure, and the resultant crude product was diluted with dichloromethane and basified with aqueous sodium hydroxide solution (1N) until pH

~ 8. The reaction mixture was extracted with dichloromethane (3x). The combined organic layers were washed with water and brine, dried over sodium sulfate, filtered, and concentrated under reduced pressure. The crude residue was purified by column chromatography (silica gel: isopropyl acetate / heptane) to give the title compound (88.4 mg, 86% yield) as a white solid.

**$^1$H NMR** (400 MHz, DMSO) *δ* 7.41 (d, J = 2.7 Hz, 1H), 7.11 (d, J = 2.7 Hz, 1H), 6.45 (d, J = 16.1 Hz, 1H), 6.16 (dd, J = 16.1, 6.9 Hz, 1H), 4.70 (s, 2H), 3.75 (s, 3H), 2.37-2.24 (m, 1H), 2.12-1.97 (m, 2H), 1.97-1.75 (m, 4H), 1.48-1.33 (m, 2H).

**$^{13}$C NMR** (101 MHz, DMSO) *δ* 152.46, 139.61, 135.47, 130.26, 124.01 (t, J = 240.6 Hz), 122.29, 121.40, 119.14, 52.81, 38.25, 32.54 (dd, J = 23.8 Hz), 28.55 (d, J = 9.8 Hz).

**HRMS (ESI+)** m/z found MH$^+$ 269.1457, $C_{14}H_{19}F_2N_2O$ requires 269.1458.

**Synthesis of C2-Acetyl**

(E)-N-(5-(2-(4,4-difluorocyclohexyl)vinyl)-6-methoxypyridin-3-yl)acetamide

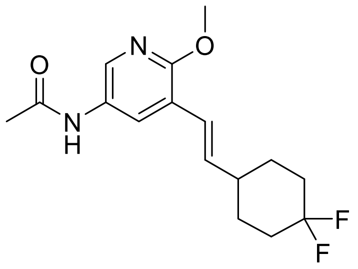


To a stirred solution of (E)-5-(2-(4,4-difluorocyclohexyl)vinyl)-6-methoxypyridin-3-amine (150 mg, 0.56 mmol) in pyridine (2.0 mL) at 15 °C under a nitrogen atmosphere was added acetic anhydride (1.0 mL) dropwise. The resulting reaction mixture was stirred at room temperature for 2 h and the reaction mixture was evaporated under reduced pressure. The crude residue was purified by column chromatography (silica gel: ethyl acetate / heptane) to give the title compound (150 mg, 86% yield) as brown solid.

**$^1$H NMR** (400 MHz, DMSO) δ 9.92 (s, 1H), 8.17 (d, J = 2.4 Hz, 1H), 7.99 (d, J = 2.5 Hz, 1H), 6.50 (dd, J = 16.3, 1.2 Hz, 1H), 6.24 (dd, J = 16.2, 6.9 Hz, 1H), 3.85 (s, 3H), 2.41-2.27 (m, 1H), 2.11-1.98 (m, 5H), 1.98-1.76 (m, 4H), 1.50-1.35 (m, 2H).

$^{13}$**C NMR** (101 MHz, DMSO) δ 168.32, 156.05, 136.84, 136.08, 130.40, 126.71, 123.98 (dd, J = 240.3 Hz), 121.74, 53.29, 38.29, 32.54 (dd, J = 23.8 Hz), 28.44 (d, J = 9.6 Hz), 23.59.

**HRMS (ESI+)** m/z found MH$^+$ 311.1568, $C_{16}H_{21}F_2N_2O_2$ requires 311.1566.

**Synthesis of C2-Methyl**

(E)-5-(2-(4,4-difluorocyclohexyl)vinyl)-6-methoxy-N-methylpyridin-3-amine

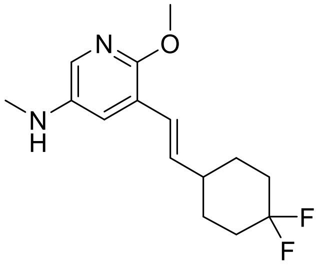


**Step 1:** Preparation of tert-butyl (E)-(5-(2-(4,4-difluorocyclohexyl)vinyl)-6-methoxypyridin-3-yl)carbamate

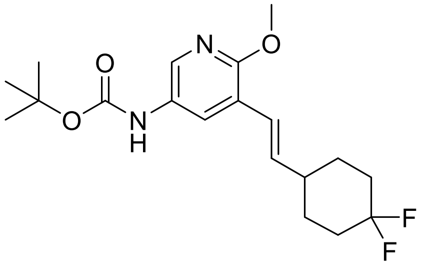


To a stirred solution of (E)-5-(2-(4,4-difluorocyclohexyl)vinyl)-6-methoxypyridin-3-amine (200 mg, 0.74 mmol) in 1,4-dioxane (10 mL) was added di-tert-butyl dicarbonate (0.21 mL, 0.89 mmol). The resulting reaction mixture was stirred at 100 °C for 3h. The cooled reaction was concentrated under reduced pressure to get crude tert-butyl (E)-(5-(2-(4,4-difluorocyclohexyl)vinyl)-6-methoxypyridin-3-yl)carbamate (250 mg, 91% yield) as brown liquid which was used directly in the next reaction step. MS (ESI+) m/z 369 (M+H)$^+$.

**Step 2:** Preparation of C2-Methyl

To a stirred solution of tert-butyl (E)-(5-(2-(4,4-difluorocyclohexyl)vinyl)-6-methoxypyridin-3-yl)carbamate (250 mg, 0.68 mmol) in tetrahydrofuran (10 mL) at 0 °C was added a solution of lithium aluminum hydride in tetrahydrofuran (1.7 mL, 3.39 mmol, 2M) under nitrogen atmosphere. The resulting reaction mixture was stirred at 70 °C for 16h. The reaction was cooled to room temperature and then quenched with ice water (2.0 mL). Aqueous sodium hydroxide solution (1N) (1.0 mL) was added dropwise, and the resulting suspension was diluted with ethyl acetate. The organic layer was washed with ethyl acetate (2 x 15 mL) and brine (3 x 10 mL). The combined organic layers were dried over sodium sulfate, filtered, and concentrated under reduced pressure. The crude was purified by column chromatography (silica gel: ethyl acetate / heptane) to give the title compound (116 mg, 61% yield) as brown viscous liquid.

**$^{1}$H NMR** (400 MHz, DMSO) *δ* 7.21 (s, 1H), 6.93 (s, 1H), 6.32 (d, J = 16.1 Hz, 1H), 6.10 (dd, J = 16.2, 6.9 Hz, 1H), 5.16-5.04 (m, 1H), 3.61 (s, 3H), 2.51 (d, J = 5.0 Hz, 3H), 2.23-2.09 (m, 1H), 1.95-1.81 (m, 2H), 1.81-1.60 (m, 4H), 1.34-1.19 (m, 2H).

**$^{13}$C NMR** (101 MHz, DMSO) *δ* 152.45, 141.44, 135.76, 127.90, 124.02 (t, J =239.6 Hz), 122.31, 119.43, 119.27, 52.85, 38.34, 32.55 (t, J = 23.3 Hz), 30.38, 28.54 (d, J = 9.0 Hz).

**HRMS (ESI+)** m/z found $MH^+$ 283.1616, $C_{15}H_{21}F_2N_2O_2$ requires 283.1617.
